# On the predictability of progression-free survival in ovarian cancer from NanoString gene expression data

**DOI:** 10.64898/2026.04.22.719856

**Authors:** Lucy B. Van Kleunen, Grace Bowman, Sarah Elizabeth Stockman, Hope A. Townsend, Logan Barrios, Kimberly R. Jordan, Rebecca J. Wolsky, Kian Behbakht, Matthew J. Sikora, Jennifer K. Richer, Junxiao Hu, Benjamin G. Bitler, Aaron Clauset

**Affiliations:** BioFrontiers Institute, University of Colorado Boulder, 3415 Colorado Avenue Boulder, CO 80303, USA; KU Leuven Campus Kulak, Department of Public Health and Primary Care, Etienne Sabbelaan 53, 8500 Kortrijk, Belgium; ITEC-imec and KU Leuven, Etienne Sabbelaan 51, 8500 Kortrijk, Belgium; Department of Ecology and Evolutionary Biology, University of Colorado Boulder, Boulder, Colorado 80302, USA; Institute for Behavioral Genetics, University of Colorado Boulder, Boulder, Colorado 80302, USA; Department of Molecular, Cellular, & Developmental Biology, University of Colorado Boulder, Boulder, Boulder, Colorado 80309, USA; Department of Computer Science, University of Colorado, Boulder, Colorado; Department of Biomedical Engineering, University of Colorado Boulder, Boulder, Boulder, Colorado 80309, USA; Department of Immunology and Microbiology, The University of Colorado Anschutz Medical Campus, Aurora, Colorado, USA; Department of Pathology, The University of Colorado Anschutz Medical Campus, Aurora, Colorado, USA; Department of OB/GYN, The University of Colorado Anschutz Medical Campus, Aurora, Colorado; Department of Pediatrics, Cancer Center Biostatistics Core, The University of Colorado Anschutz Medical Campus, Aurora, Colorado; Santa Fe Institute, Santa Fe, New Mexico

## Abstract

In the treatment of high grade serous ovarian cancer (HGSC), patients initially diagnosed with unresectable tumors are first treated with neoadjuvant chemotherapy (NACT) to reduce tumor burden prior to surgery. Analysis of matched pre- and post-NACT samples from the same patients enables the investigation of chemotherapy impacts and the biomarkers of progression. Although the tumor immune microenvironment (TIME) has increasingly been recognized as critical in shaping the development and progression of HGSC, we lack a comprehensive understanding of how chemotherapy remodels the TIME. Previous studies have found evidence for a general inflammatory response post-NACT, despite inconsistencies regarding which differentially expressed genes and pathways are implicated. We combine matched NanoString gene expression data from multiple sources to create a large dataset of matched pre- and post- NACT samples (N=83, with 29 novel to this study) and investigate reproducibility. Further, we use machine learning methods to investigate whether patient progression-free survival (PFS) can be predicted from the observed impact of chemotherapy on the TIME as represented by the comprehensive set of NanoString features. We find overall low predictability of PFS from all NanoString features, suggesting that previous results may have been limited by small sample size effects and that larger datasets are needed to identify more generalizable and translatable findings. We identify a set of differential expression features that are the most important for predicting patient outcomes that can be validated in future computational and biological studies.

**Author summary:** A subset of patients with high grade serous ovarian cancer are treated with chemotherapy before surgery to reduce tumor burden. We investigate a large dataset of samples taken before and after chemotherapy. These matched samples enable an investigation of how the environment around tumors, for example immune cell infiltration, reacts to chemotherapy, providing insights into biomarkers for treatment response and treatments that could complement chemotherapy. This larger dataset only partially replicates results from previous studies, while also providing new insights. Machine learning models designed to predict the time to patient recurrence from available biomarkers indicate that they are not strongly predictive of patient outcomes, in contrast to past studies. These results suggest that larger datasets are needed. We identify a set of genes that change with chemotherapy and are indicative of and potentially useful for predicting time to disease recurrence and can be further investigated.

## Introduction

High grade serous ovarian carcinoma (HGSC) of the ovary and fallopian tube is the most common histotype of ovarian cancer (Bowtell 2010, Shih et al. 2021). This cancer is distinguished by late-stage diagnosis, acquired therapy resistance, and nearly inevitable disease recurrence, motivating research into new treatments (Siegel et al. 2025). A typical course of treatment involves surgical cytoreduction followed by platinum/taxane-based chemotherapies. Prior to surgical cytoreduction, a laparoscopic biopsy sample is taken to diagnose malignancy and tumor type, after which tumor resectability is assessed. Patients with unresectable tumors are first treated with neoadjuvant chemotherapy (NACT) in order to shrink the tumor before further treatment (Fagotti et al. 2020). In this study, we build on prior research taking advantage of matched biopsy (pre-NACT) and surgical (post-NACT) samples to study how chemotherapy remodels the tumor immune microenvironment (TIME) in HGSC, and the predictability of patient outcomes from standard TIME biomarkers.

In a previous study (Jordan et al. (2020)), we showed that NACT induced the activation of several oncogenic pathways and identified 18 genes whose differential expression pre- and post-NACT correlated with progression-free survival (PFS). Since then, several studies have investigated how differential gene expression and TIME remodeling relate to treatment response and prognosis. James et al. (2022) reported upregulation of immunoregulatory and inflammatory pathways following NACT, supporting the association between post-NACT inflammatory activation and shorter PFS in Jordan et al. These findings indicate a possible shift toward a more immunosuppressive microenvironment following chemotherapy, given an upregulation of immune inhibitory molecules. Zhang et al. (2022) similarly identified enhanced expression of genes involved in stress response, many of which are implicated in inflammatory and immune-modulatory processes. For example, stress responses mediated through *CEBPB* can promote inflammatory responses following chemotherapy (Zhang et al. 2022). Overall, these studies have identified several interconnected genes, including *SGK1*, *IL6*, and *DUSP1*, which are involved in stress response and immune signaling pathways and may be potential targets for therapies that could be used in conjunction with chemotherapy to extend PFS.

However, other studies have suggested that the role of the immune system and inflammation may be context dependent. McAdams et al. (2024) focused specifically on mast cells, reporting increased mast cell abundance and histamine expression (a marker of mast cell degranulation) post-NACT. Lower levels of pre-NACT mast cells were significantly associated with improved PFS, implying that mast cells may play an immunosuppressive role, even before treatment. Hence, certain kinds of immune cell infiltration (such as infiltration by mast cells) may lead to worse outcomes. Other studies have found that immune cell infiltration is correlated with better patient outcomes. For example, Barna et al. (2023) found that patients with tumors with greater levels of immune cell infiltration (especially CD4+ T cells) had higher overall survival, and Lee et al. (2020) found that patients who had no visible disease after gross resection had a higher number of infiltrated T cells in the resected tumor as compared to patients who required NACT. And, Denisenko et al. (2024) used spatial transcriptomics to find evidence of discrete subclonal zones of different inflammatory signals within the tumor microenvironment, suggesting that the complexity of the TIME can confound efforts to identify general mechanisms. This literature reinforces the importance of the TIME for understanding HGSC progression and response to treatment. However, it also highlights inconsistencies regarding which differentially expressed genes and pathways are implicated, likely driven in part by heterogeneity in spatial organization and cell type composition, and underscoring the complexity of translating gene expression signals to actionable biomarkers or targets in rare diseases.

A key limitation of past studies on matched pre- and post-chemotherapy samples is their limited samples size. Here, we combine matched NanoString gene expression data from 4 sources to create a large dataset of matched pre- and post- NACT samples (N=83, with 29 novel to this study). We investigate whether prior single-cohort results can be replicated in this multi-cohort dataset, and we use machine learning methods to predict PFS to identify key NanoString features that could act as predictive biomarkers. Toward this end, we conduct a systematic analysis to investigate whether results from Jordan et al. 2020 identifying 18 genes as biomarkers replicate in this larger multi-cohort dataset, as well as precisely isolate which changes to the analysis, if any, drive any analytic differences. Specifically, within a 2x2x2 design, we vary (1) the dataset size (Jordan et al. N=6 dataset vs. the expanded N=83 multi-cohort dataset including all Jordan et al. samples as a subset), (2) sample normalization vs. un-normalized sample values, and (3) statistical or machine learning methodology. While standard normalization procedures are typically applied to NanoString gene expression data (see Materials & Methods), we chose to include normalization as an axis in our analysis because normalization is performed across entire cohorts. Thus, feature values for the same samples between normalized single-cohort datasets used in prior work and the normalized multi-cohort dataset can differ. Considering raw values removes these whole-cohort effects. As neither approach was clearly superior, we considered both in our analysis to investigate the impact on biomarker identification. We find that our results from a larger dataset partially replicate prior work, but that models based on all NanoString PanCancer IO 360 gene expression, cell type, and signaling features show poor predictive performance for PFS. Despite low predictive performance, we identify a set of top features which are the most important for predicting PFS in multi-dimensional models. These features, including NACT-induced changes to *MTOR* and *RAD51*, can be investigated in future work as potential biomarkers in HGSC.

## Results

### Differences pre- and post- NACT

First, we characterized the differences in NanoString biomarkers between pre-and post- NACT samples in our large multi-cohort dataset. This descriptive analysis provided a baseline for interpreting our subsequent replication analysis of the Jordan et al. results based on patient-level log-fold change features by characterizing the overall impact of NACT on the TIME for both normalized and non-normalized versions of the dataset. It also validated that our multi-cohort dataset showed changes post-NACT supported by prior studies on ovarian cancer biology, supporting its use for identification of generalizable biomarkers.

The multi-cohort dataset with 83 patients showed a median patient age at diagnosis of 63 years and a median PFS of 10 months (Fig. 1A,B). To examine NACT-induced global changes to the TIME, we compared 750 gene expression, 13 cell type, and 25 signaling features pre- and post- NACT. We also compared results on normalized vs. non-normalized versions of the dataset (see Materials & Methods). We found 414 features to have a significant change post-NACT based on paired t-tests in the normalized data and 365 in the non-normalized data. With a multiple testing correction applied (Benjamini-Hochberg method), 360 remained significant for the normalized data, and 305 remained significant for the non-normalized data, indicating a significant global change to the TIME post-NACT. The full list of post-NACT differential expression results is available in Supplemental Table 1.

**Figure 1.**
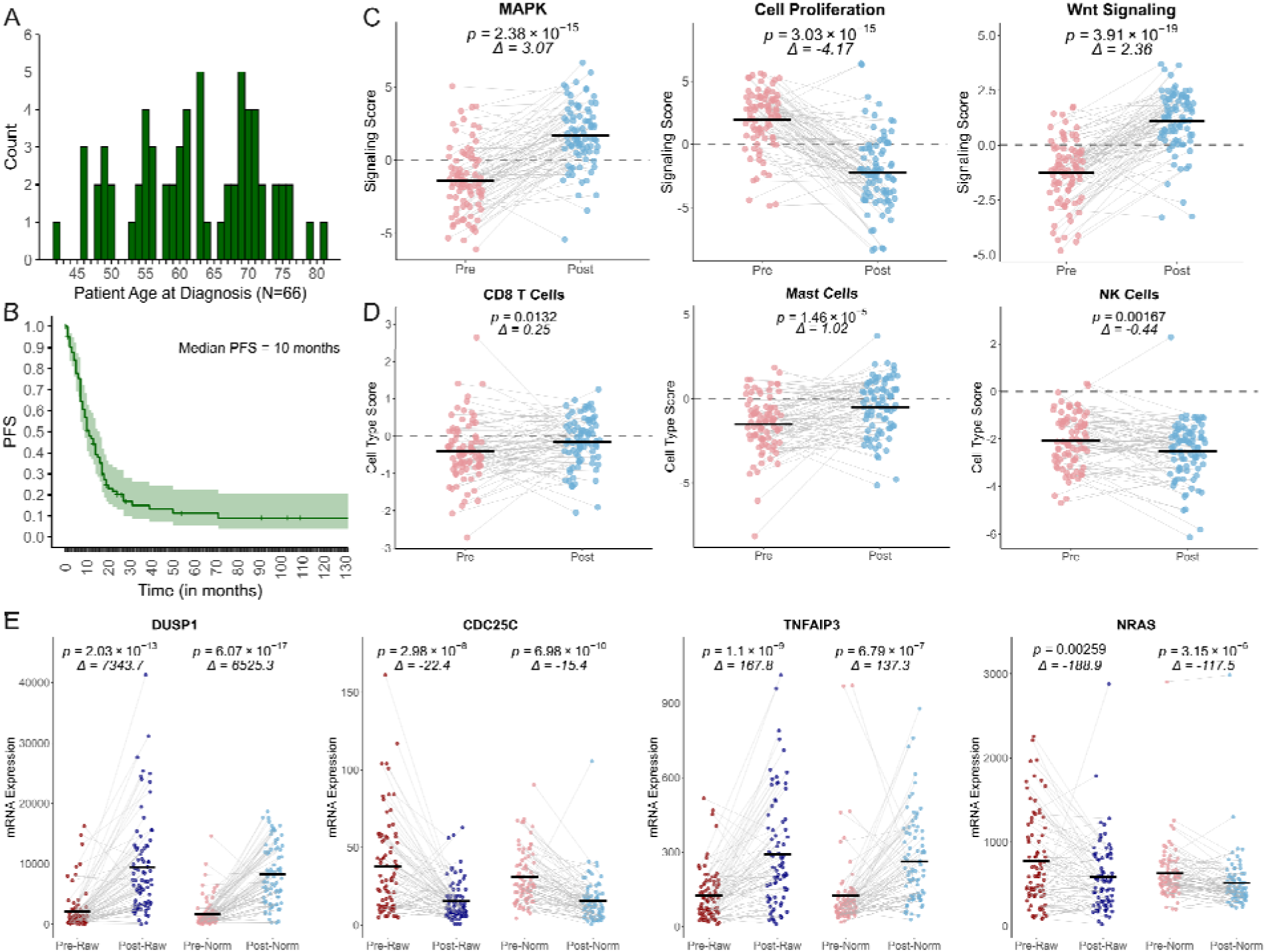
Characteristics of the N=83 dataset of pre- and post- NACT matched samples. (A) Histogram of patient age at diagnosis for the N=66 matched samples for which this information is available. (B) Kaplan-Meier survival curve for PFS showing recurrence probability over time. (C) Select results for signaling feature differences pre- and post-NACT for normalized data. (D) Select results for cell type feature differences pre- and post- NACT for normalized data. (E) Select results for gene differential expression pre-and post- NACT, for non-normalized (Pre/Post-Raw) and normalized (Pre/Post-Norm) data. For plots C-E, deltas show the mean difference pre- and post- NACT and p-values are shown for paired t-tests, not adjusted for multiple comparisons.

Amongst significant cell type and signaling features, we examined a subset of these features based on prior work indicating their importance in ovarian cancer biology (MAPK signaling: Hew et al. 2016, cell proliferation: Mahadevappa et al. 2017, Wnt signaling: Bishara et al. 2025, CD 8+ T cells: Goode et al. 2017, Mast cells: Cao et al. 2021, NK cells: Henriksen et al. 2020). We identified an increase in MAPK signaling (normalized: *t*=-9.74, *p*=2.38e-15), decrease in cell proliferation (normalized: *t*=9.69, *p*=3.03e-15), increase in Wnt signaling (normalized: *t*=-11.68, *p*=3.91e-19), increase in CD8+ T cells (normalized: *t*=-2.53, *p*=0.0132), increase in mast cells (normalized: *t*=-4.61, *p*=1.46e-5), and decrease in NK cells post-NACT (normalized: *t*=3.25, *p*=0.00167) (Fig 1C,D). These results are consistent with previous studies in which CD8+ T cells (Lodewijk et al. 2022) and mast cells (McAdams et al. 2024 and Lodewijk et al. 2022) were upregulated post-NACT, while James et al. demonstrated post-NACT increases in MAPK and Wnt signaling and a decrease in cell proliferation.

Given our focus on replication of prior single cohort results on a larger multi-cohort dataset, we specifically examined the 12 cell-type and signaling features that were found in Lodewijk et al. 2022 to increase post-NACT for patients with no or partial tumor response to NACT, representing an inflammatory signature. We investigated the change in these features post- NACT on the multi-cohort dataset reduced to patients with worse prognosis (PFS < 12 months, N=42 patients). Out of the 12 inflammatory signature features, we found that 8 showed a significant increase post-NACT in the normalized data for this population (Antigen Presentation *t*=-2.50 *p*=0.016, Costimulatory signaling *t*=-2.75 *p*=0.009, Cytokine and Chemokine Signaling *t*=-3.58 *p*=0.0009, Immune Cell Adhesion and Migration *t*=-2.51 *p*=0.016, Lymphoid Compartment *t*=-2.73 *p*=0.0093, Myeloid Compartment *t*=-2.15 *p*=0.0373, Total TILs *t*=- 2.60 *p*=0.0128, and T cells *t*=-2.47 *p*=0.0175). The same features showed a significant increase post-NACT in the non-normalized data, except for T cells, which did not show a significant change. Macrophages showed a significant decrease post-NACT in the normalized data (*t*=2.59, *p*=0.0133), and no significant change in the non-normalized data. In both cases we considered non-adjusted p values (Fig S2).

Amongst our gene expression results on the entire multi-cohort dataset (Fig. S1, Supplemental Table 1), we specifically investigated a subset of gene expression features known to be relevant to ovarian cancer biology based on prior work (*DUSP1*: Sanders et al. 2022; *CDC25C*: Fei et al. 2019; *TNFAIP3*: Hee Han et al. 2021; *NRAS*: Zhong et al. 2019), seeing an increase in *DUSP1* expression in line with results in James et al. and Jordan et al. (normalized: *t*=-10.55, *p*=6.07e-17; non-normalized: *t*=-8.77, *p*=2.03e-13), decrease in *CDC25C* expression (normalized: *t*=6.98, *p*=6.98e-10; non-normalized: *t*=6.13, *p*=2.98e-8), increase in *TNFAIP3* expression (normalized: *t*=-5.38, *p*=6.79e-7, non-normalized: *t*=-6.88, *p*=1.10e-9), and decrease in *NRAS* expression post NACT (normalized: *t*=5.00, *p*=3.15e-6, non-normalized: *t*=3.11, *p*=0.000259) (Fig. 1E).

### Statistical analyses only partially replicate prior results on whole cohort

In order to isolate the analytic effects of changing the data set size (N=6 vs. N=83), whether the biomarker values are normalized or not, and whether we use standard statistical methods or machine learning methods, we conduct a systematic analysis of all pairs of changes along these three dimensions. In this systematic 2x2x2 evaluation design, the first analysis replicates the statistical analyses connecting pre-and post- matched features to PFS performed in prior work (Jordan et al. 2020). This formal replication provides a baseline for cleanly interpreting the results from the other variations in our design. In these analyses, we transformed 788 gene expression, cell type, and signaling features into log-fold change features for each of the 83 matched samples.

The baseline analysis exactly replicated the univariate Pearson correlation analysis from Jordan et al. 2020 on the 750 gene expression log-fold change features from the original set of 6 samples. We recovered the same 18 genes with log-fold change values correlated to PFS as in the original work (Fig. 2A). Notably, when normalization was based on the full N=83 multi-cohort data set, rather than on the smaller N=6 data set, this same analysis of the 6 samples only partially replicated prior results (Fig. S3). If the biomarker values for the N=6 samples were not normalized at all, again, prior results only partially replicate (for 5/18 features; Fig 2B, results shown across all 788 features). These differences highlight the sensitivity of the statistical results to normalization and the importance of reporting normalization constants with results (see normalization constants for the N=6 and N=83 data, Supplemental Table 2, 3). We performed this same analysis on the larger N=83 dataset and analyzed Pearson correlations between these 788 log-fold change features and PFS. We found 122 features significantly correlated with PFS for the normalized version of the dataset and 95 features significantly correlated for the non-normalized version (full results are listed in Supplemental Table 4). On this larger and more diverse data set, only one of the originally identified features from Jordan et al. 2020, *CCNO* (r= 0.903, p=0.014 in Jordan et al. 2020) was found to correlate significantly with PFS in the multi-cohort dataset (normalized: r= -0.248, p=0.024; non-normalized: r=-0.275, p=0.012), but with the opposite direction of effect.

**Figure 2.**
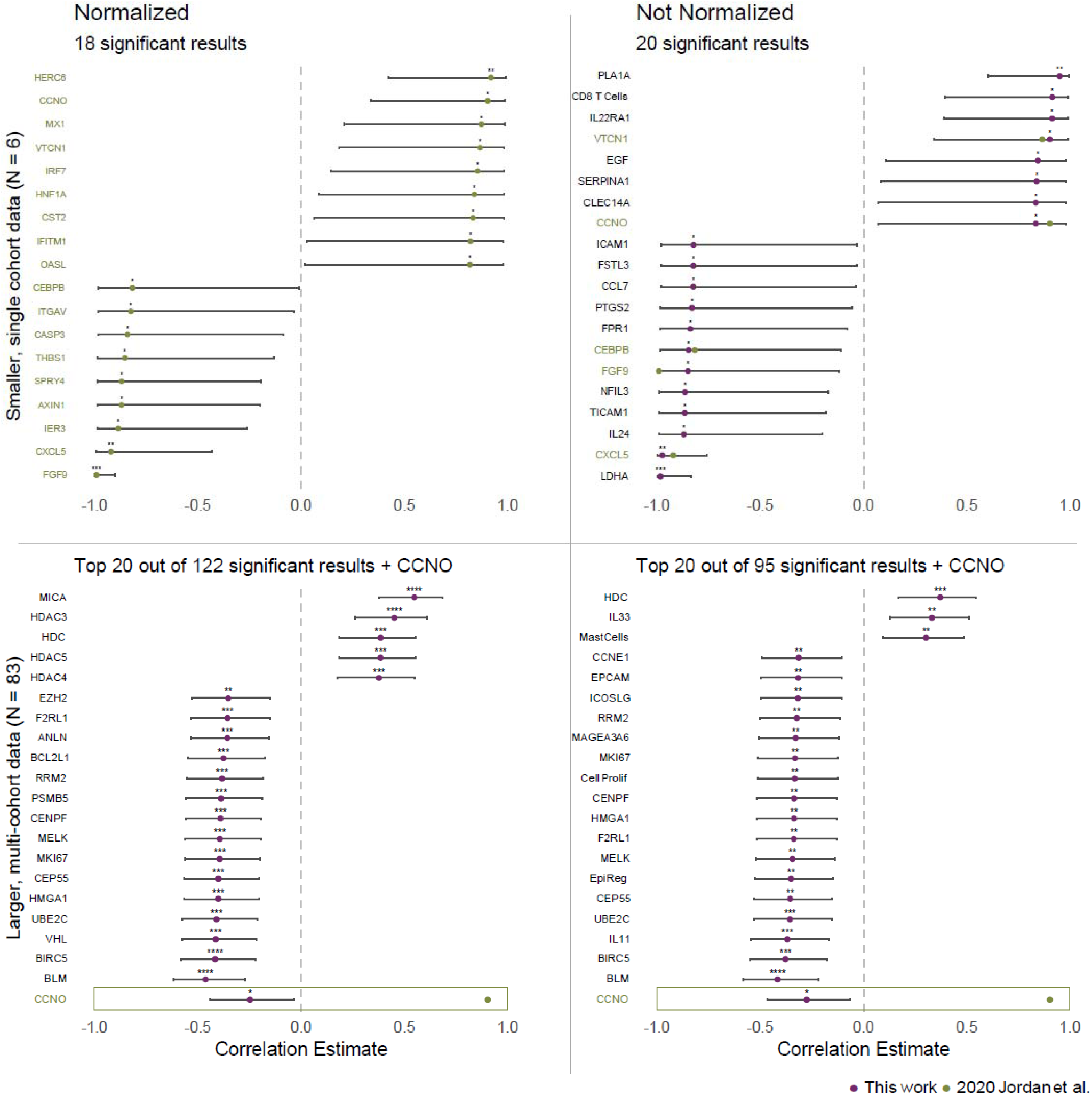
Pearson correlation results between log-fold change features and PFS. (A) Significant results from the analysis run on the N=6 samples from Jordan et al. 2020, with the dataset normalized across the N=6 samples, for 750 gene features. (B) Significant results from the analysis run on the N=6 samples from Jordan et al. 2020, non-normalized. (C) Top 20 out of 122 significant results from the analysis run on the N=83 normalized samples. (D) Top 20 out of 95 significant results from the analysis run on the N=83 non-normalized samples. Analyses for B-D were run over 788 gene, cell, and signaling features. Plots show Pearson correlation estimates and confidence intervals, with a vertical line at 0, asterisks indicate significance levels (non-adjusted, **** p< 0.0001, *** p< 0.001, ** p<0.01, *p<0.05). Top 20 results were determined by ranking based on the absolute value of the correlation estimates. Green dots indicate estimates for features in the original Jordan et al. 2020 analysis and feature labels are colored green if they were significant in the Jordan et al. 2020 results. Plots C and D show *CCNO* along with the top 20 results as this was the only feature significant in the N=6 analysis that was still significant in the N=83 analysis, though it switched directionality.

These results were also sensitive to normalization (Fig 2C,D). After performing a multiple testing correction (Benjamini-Hochberg method), 27 features remained significant for the normalized data on the multi-cohort dataset (negative estimates: *ANLN* r=-0.357, p=0.039; *BCL2L1* r=-0.377, p=0.021; *BIRC5* r=-0.416, p=0.016; *BLM* r=-0.463, p=0.004; *CCNE1* r=-0.340, p=0.048; *CENPF* r=-0.390, p=0.018; *CEP55* r=-0.401, p=0.017; *CRABP2* r=-0.341, p=0.048; *ENO1* r=-0.341, p=0.048*; EPCAM* r=- 0.350, p=0.406*; EZH2* r=-0.353, r=0.040; *F2RL1* r=-0.357, p=0.039; *HMGA1* r=-0.401, p=0.017; *MELK* r=-0.394, p=0.018; *MKI67* r=-0.394, p=0.018; *PSMB5* r=-0.388, p=0.018; *RRM2* r=-0.383, p=0.018; *UBE2C* r=-0.409, p=0.016; *VHL* r=-0.413, p=0.016; Epigenetic Regulation r=-0.350, p=0.041; positive estimates: *HDAC3* r=0.452, p=0.005; *HDAC4* r=0.376 p=0.021; *HDAC5* r=0.384, p=0.018; *HDC* r=0.385, p=0.018; *IL33* r=0.353, p=0.040; *MICA* r=0.547, p=6.71e-5; *RBL2* r=0.343, p=0.048; see Supplemental Table 4). All adjusted p values were significant at the level of p<0.05, while *HDAC3* and *BLM* were significant at the p<0.01 level, and *MICA* was significant at the p<0.0001 level. No features remained significant after multiple testing correction for the non-normalized multi-cohort dataset.

As performed in James et al. 2022, we performed univariate Cox regressions between pre- and post-NACT cell type and signaling features (Fig S4). We found largely the same results for the original N=31 samples from James et al. (13 out of 18 significant results), with differences attributable to our normalization over N=83 samples (Fig S4A). In the larger, multi-cohort data set (N=83), we found that 10 out of the 18 significant results from James et al. 2022 were reproduced for the normalized and non-normalized data (Fig. S4C,D). Considering non-adjusted p values on the N=83 dataset, the only features amongst this set that were significant at the p<0.0001 level for both the normalized and non-normalized datasets were post-NACT autophagy, with a hazard ratio estimate < 1 and post-NACT cell proliferation, with a hazard ratio estimate > 1.

### Machine learning analyses based on all NanoString features show poor predictive performance on the whole cohort

Machine learning tools represent a powerful analytic method for characterizing potentially complicated correlations between numeric features like patient biomarker values and outcome variables like PFS. Of particular value is their reliance on out-of-sample “testing” or cross-validation to measure accuracy, i.e., model parameters are estimated using a subset of the data (training data) and model quality is evaluated on “held out” data that the model has not seen before (test data), and in some cases, their ability to learn non-linear relationships between dependent and independent variables. In the final branch of our systematic evaluation, we assess whether changing from standard statistical methods to modern predictive modeling methods can uncover statistically valid biomarker relationships with patient outcome. Specifically, we tested the ability of several machine learning algorithms to learn to predict out-of-sample PFS based on all 788 of the log-fold change features considered together. We treated this task as a binary prediction problem by transforming PFS into a binary outcome split at median PFS (10 months, see Materials & Methods). We tested random forests, which can learn non-linear relationships, and logistic regression with three different penalty terms (L1, L2, and Elastic-Net).

We observed high ROC-AUC predictive performance on the smaller Jordan 2020 dataset. However, the sample size was much smaller (N=6) than sample sizes typically investigated for machine learning models and the observed performance did not always exceed the distribution of randomized feature permutation tests (Fig. 3A,B).

**Figure 3.**
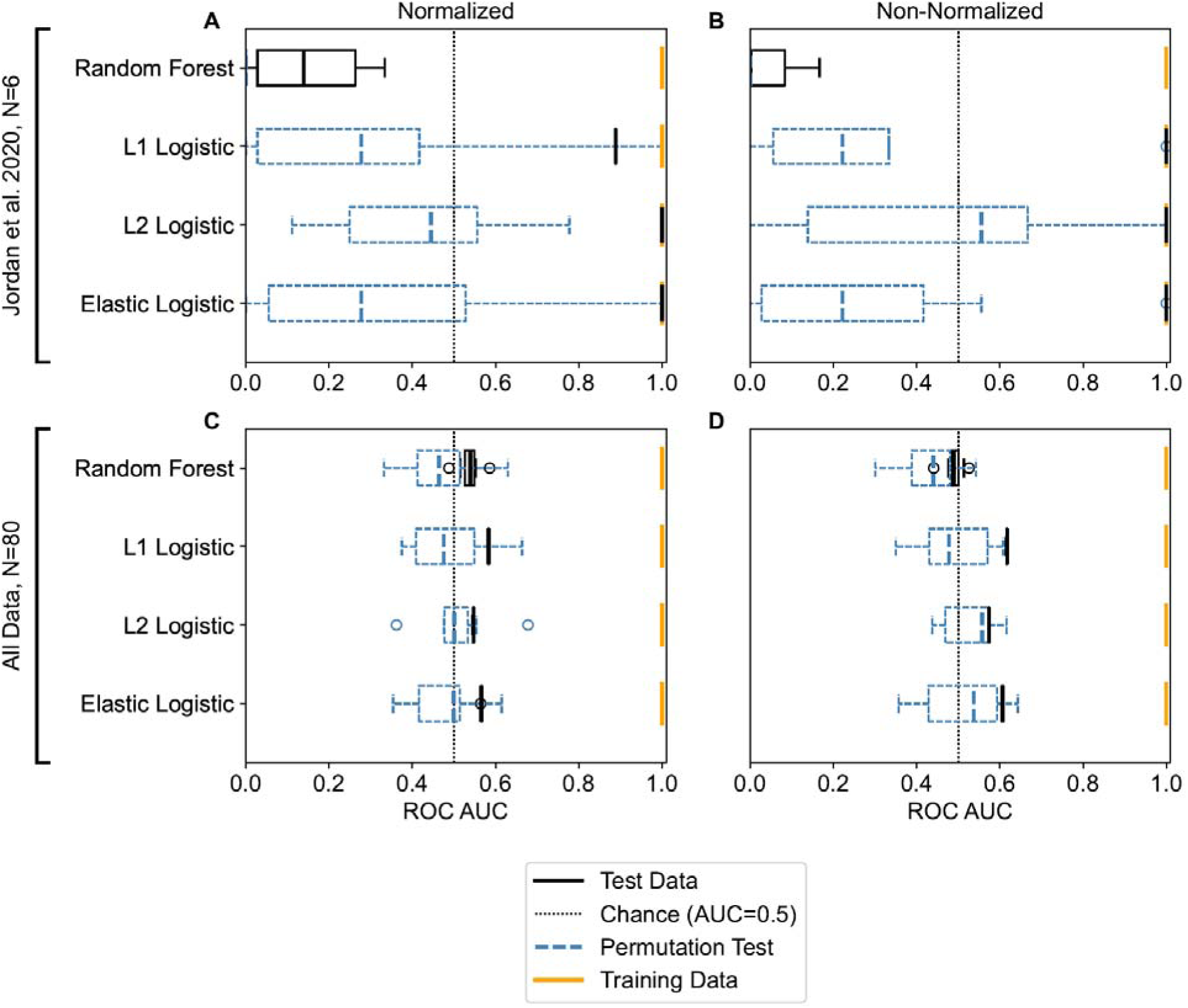
Predictive performance results for binary prediction of PFS based on the ROC-AUC metric across 10 iterations of leave-one-out cross-validation. (A) For the N=6 samples from Jordan et al. 2020 on the normalized dataset, (B) for the N=6 samples from Jordan et al. 2020, non-normalized, (C) for the N=80 samples, normalized, and (D) for the N=80 samples, non-normalized. Normalized versions of the data are normalized over all N=83 samples in the dataset. The x axis of the plot ranges from 0 to 1 based on the range of possible ROC-AUC values, and a vertical dashed line is shown at ROC-AUC=0.5, which represents a baseline value above which predictive performance is better than a random guess. The distribution of results on the test dataset across the iterations is shown in black. The mean performance on the training dataset across iterations is shown in yellow. The distribution of results from dataset-specific feature permutation tests are shown in blue (dashed). Box plots show the median and the whiskers represent 1.5 times the interquartile range.

On the larger dataset (N=80, after excluding the 3 samples censored before the median) we saw performance that was equivalent or only slightly higher than the ROC-AUC baseline performance of 0.5 (Ling et al. 2003), as well as results that mostly fell within the distribution of dataset specific baseline tests with randomized features, indicating either no or a very weak predictive signal for all features considered together, as well as model overfitting based on perfect performance on the training dataset (Fig. 3C,D). Evaluating over 10 iterations of leave-one-out cross-validation, we saw the best mean performance on the N=80 dataset for L1 Logistic regression for both the non-normalized (mean AUC=0.62±0) and the normalized (mean AUC= 0.58±0) datasets.

Other metrics of predictive performance are in Supplemental Table 5. Predicting within each of the other 3 sources, predictive performance was either equal to or better than that on the N=80 dataset for the best models (Fig. S5). Several variations in the analysis did not improve out-of-sample performance.

For instance, we did not find better predictive performance for the N=80 dataset if treating the prediction task as multi-class prediction of chemotherapy status (see Materials & Methods) (Fig. S6). We did not see better predictive performance for the N=80 dataset for logistic regression with optimization of the regularization strength hyperparameter (Supplemental Table 6), or for the random forest with tests of different hyperparameter combinations (Fig. S7). And, binary prediction results on the N=80 dataset were also not better without feature scaling before prediction (Fig. S8) and looked similar with Age included as an additional feature (Fig. S9).

Moreover, two variations in the machine learning algorithms failed to improve out-of-sample performance. These sensitivity tests assessed whether poor predictive performance was due to treating the problem as a binary prediction rather than as a survival analysis task. We predicted PFS directly out-of-sample taking censoring into account across the entire N=83 dataset using two common machine learning approaches: random survival forests and regularized Cox proportional hazards models, measuring performance in this case using Harrell’s C-index. For both random survival forest (mean C-index=0.608±0.066 on non-normalized and mean C-index=0.519±0.006 on normalized) and the regularized Cox proportional hazards model (mean C-index=0.556±0.086 on non-normalized data, mean C-index=0.544±0.039 on normalized data), we found performance results only slightly higher than baseline, in line with our binary prediction results.

Finally, we performed a sensitivity test to assess whether feature values clustered according to clear differences by data source---a pattern that can be exploited by machine learning algorithms to predict cohort-correlated recurrence patterns rather than predict based on the underlying biology (as in "shortcut learning" in medical imaging prediction tasks, Banerjee et al. 2023). We trained multi-class machine learning models to predict which of the 4 data sets a patient record came from, using only the 788 features. We found predictive performance only slightly better than chance AUC > 0.5 but in a similar range as the predictive performance results for binary prediction of PFS. This concurrence indicates a marginally detectable difference per source (Fig S10). However, a visual analysis of the samples embedded in their first 2 principal components showed no obvious clustering by source or binarized progression-free survival (Fig. 4).

**Figure 4.**
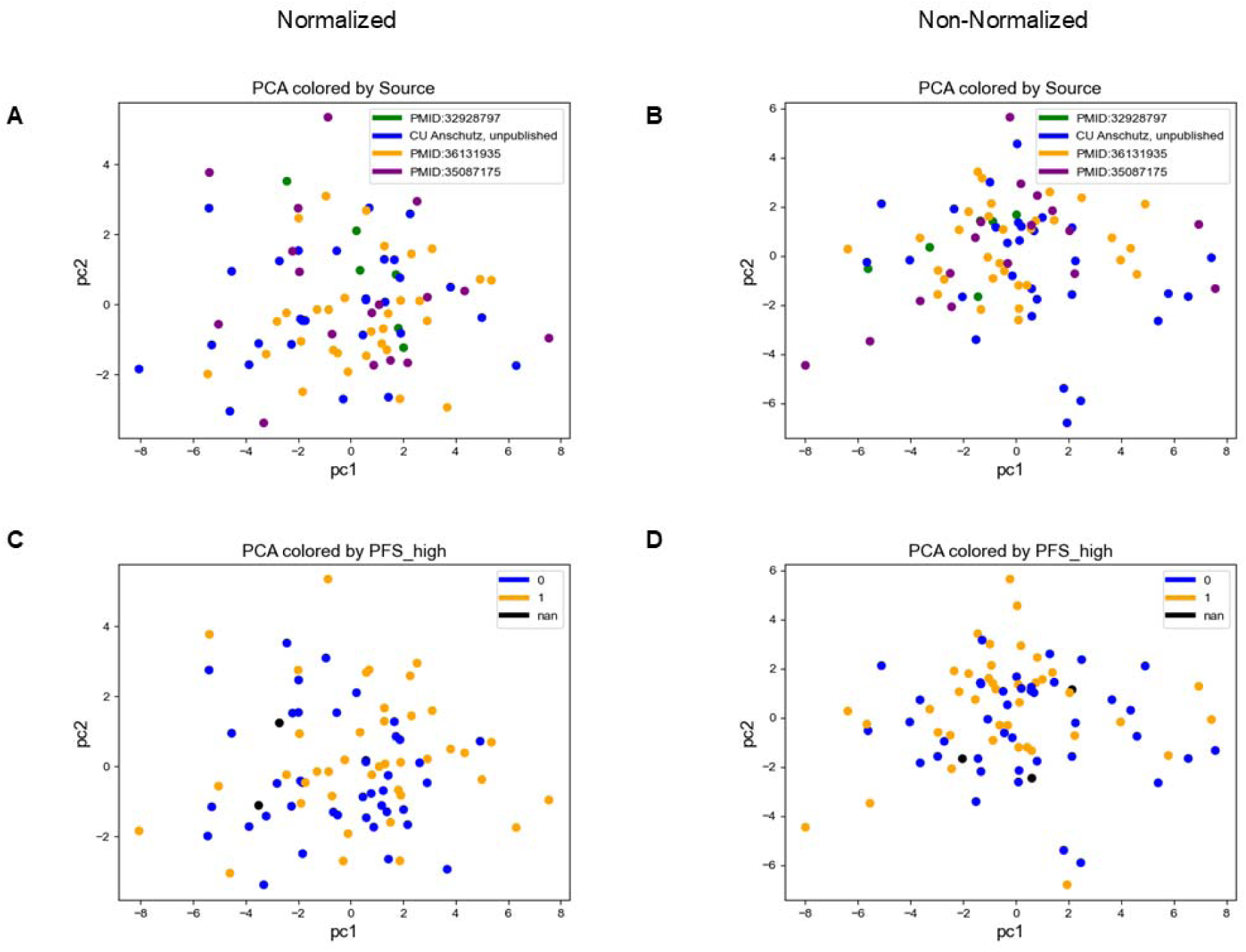
A visualization of all N=83 samples on the 2D axis of the first two principal components (pc1, pc2) determined via principal component analysis. (A) Normalized data, colored by data source. (B) Non-normalized data, colored by data source. (C) Normalized data, colored by the binary progression-free survival outcome. (D) Non-Normalized data, colored by the binary progression-free survival outcome.

### Evaluating feature importance

Despite low predictive performance and overfitting for models built from all NanoString features considered together to predict PFS, examining the most important features used to build these multidimensional models can be used as an alternative strategy for biomarker identification that also takes into account how features impact performance in combination with other features. These results can then be compared to multi-cohort results for biomarker identification based on univariate statistical analyses which consider features individually.

To identify the most important features driving predictive performance, we considered the ranking of feature importances from the 8 binary PFS classification models trained on the normalized and non-normalized data for the N=80 dataset (Fig S11, S12). We combined these rankings using the Borda method to produce an aggregated list of top features (Fig. 5).

**Figure 5.**
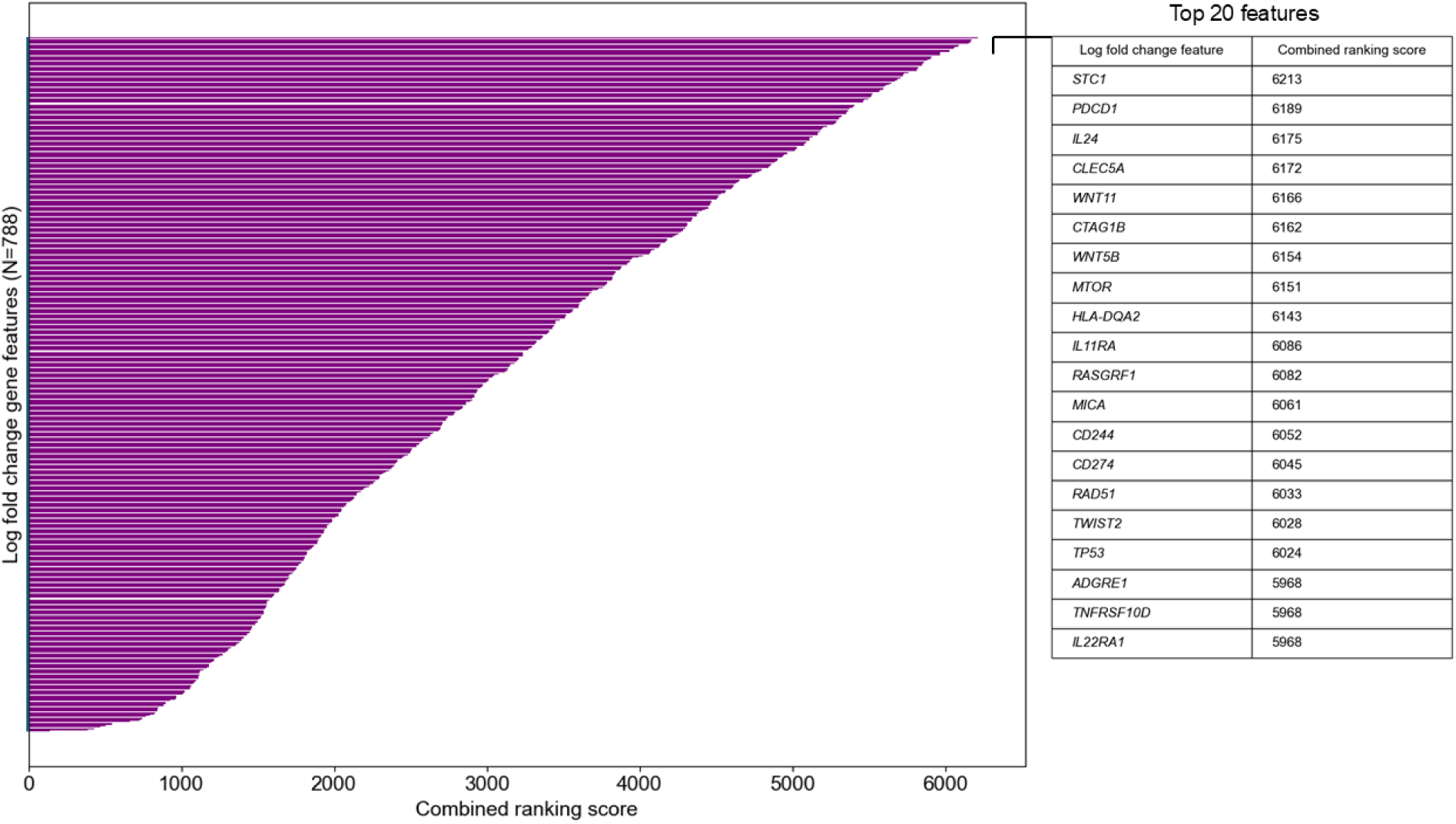
Top 20 log fold change features identified via rank aggregation. The top 20 features were determined by combining the rankings provided by 8 predictive models using the Borda method, which produces a combined ranking score for each feature. The combined ranking scores across all 788 features are shown with the top 20 highlighted.

As a sensitivity test, we also examined the most important log fold change features identified from a regularized multidimensional Cox proportional hazards model fit on the entire dataset. This model kept 106/788 features with non-zero coefficient values for the normalized and 173/788 for the non-normalized. It identified 8 out of the same top 20 features (*MICA*, *WNT5B*, *STC1*, *PDCD1*, *TNFRS10D*, *HLA-DQA2*, *MTOR*, *CTAG1B*) in the normalized dataset and 10 out of the top 20 (*ADGRE1*, *CTAG1B*, *PDCD1*, *WNT5B*, *STC1*, *TNFRSF10D*, *MICA*, *WNT11*, *CD274*, *TP53*) in the non-normalized dataset (Fig S13).

We looked at which of these top 20 log fold change features were also significant in the Pearson correlations on the multi-cohort dataset based on non-adjusted p values (5/20 for non-normalized: *CLEC5A* r=-0.269 p=0.014; *IL11RA* r=0.219 p=0.046; *IL22RA1* r=-0.227 p=0.039; *MICA* r=0.277 p=0.011; *RAD51* r=-0.264 p=0.016; 5/20 for normalized: *CD274* r=0.222 p=0.046*; CLEC5A* r=-0.229, p=0.037; *IL11RA* r=0.306 p=0.005; *MICA* r=0.547 p=8.52e-8; *RAD51* r=-0.277 p=0.011) (Fig S14, Fig S15), and which had significant log-rank tests on the multi-cohort dataset if splitting Kaplan-Meier curves between low (<=median) and high (>median) log fold change values, which is another univariate statistical approach often used to identify biomarkers in ovarian cancer. We found 7/20 for non-normalized: *CLEC5A* p=0.016, *CTAG1B* p=0.042, *IL11RA* p=0.044, *IL22RA1* p=0.038, *MTOR* p=0.017, *RAD51* p=0.03, *TP53* p=0.044 and 6/20 for normalized: *CLEC5A* p=0.014, *IL11RA* p=0.031, *MICA* p=0.038, *MTOR* p=0.005, *RAD51* p=0.021, *TNFRSF10D* p=0.019 (Fig S16, S17). These results indicate that considering features univariately in their relationship to PFS led to somewhat different results than considering them in multidimensional models.

We also examined the global shift pre- and post- NACT for these top 20 identified genes to better understand their relationship to overall TIME remodeling, finding that 8 of these genes had a significant shift pre- and post-chemotherapy for either the normalized or non-normalized version of the data (up-regulated post: *IL11RA t*=-3.92, *p*=0.0002; *MICA t*=-2.78*, p*=0.0067; *CD274 t*=-2.13, *p*=0.0356; *CD244 t*=-2.37, *p*=0.0200; *TNFRSF10D t*=-4.01, *p*=0.0001; down-regulated post: *WNT11 t*=1.97, *p*=0.0521; *MTOR t*=4.16, *p*=7.81e-5; *RAD51 t*=4.03, *p*=0.0001; Fig. S18).

## Discussion

The remodeling effect of chemotherapy on the tumor microenvironment (TIME) in high grade serous ovarian cancer (HGSC) has been strongly implicated in differential patient responses to chemotherapy, and eventual chemo-resistance. A better understanding of, for example, the role of inflammatory responses or immune infiltration post-NACT, would inform efforts to develop new therapies to complement NACT to improve patient survival. However, inconsistencies across past studies have complicated efforts to identify which differentially expressed genes and pathways are implicated. We carried out a systematic analysis of NanoString gene expression data in matched pre- and post- NACT samples to assess the degree to which prior results were due to differences in sample size, composition, and biomarker normalization, and we assessed the impact of the choice of analytical approach, specifically contrasting more standard univariate statistical analyses with modern multidimensional machine learning models. Under this systematic 2x2x2 design, while we found evidence for a global change in post-NACT samples in line with previous work, we did not replicate previous results for which particular log fold change features explained a difference in PFS in our large, multi-cohort dataset. Specifically, results for the 18 gene biomarkers whose differential expression pre- and post-NACT correlated with PFS in Jordan et al. (2020) did not replicate in a larger multi-cohort dataset based on either statistical or machine learning analyses using either normalized or non-normalized versions of the dataset.

The dataset in Jordan et al. 2020 was small (N=6) and possibly was less representative of the diversity in HGSC presentation than the N=83 multi-cohort dataset. We also found poor out-of-sample predictive performance for all NanoString features considered together on the multi-cohort dataset, only slightly beating baseline (random) performance. We identified a new set of log fold change features that were important for predicting patient prognosis in this dataset that could be further investigated, including in machine learning models built from only these biomarkers, which might show reduced overfitting and higher out-of-sample predictive performance.

In our study, we observed that a higher *MTOR* log fold change following chemotherapy is associated with worse PFS (Fig. S16, S17) and that this was an important feature for predicting PFS in multidimensional models (Fig. 5). *MTOR* is a central regulator of multiple downstream outputs including protein synthesis, metabolism, cell survival, autophagy, and cytoskeletal dynamics (reviewed in Panwar et al. 2023). *MTOR* activity is dependent on the association with specific protein complexes, notably mTORC1 (Raptor-associated) and mTORC2 (Rictor-associated).

While mTOR-related signaling can drive chemotherapy resistance, stemness, and immune remodeling (Pi et al. 2021, Chang et al. 2013, Zhou et al. 2007, Deng et al. 2019), single agent mTOR inhibitors have had limited anti-tumor responses (Emons et al. 2016, Van Der Ploeg et al. 2021). Additionally, there are reports that mTOR inhibition can drive the stemness in certain cancer contexts (Yang et al. 2011). Interestingly, a combination of mTOR inhibition with PI3K inhibitor or chemotherapy enhances an anti-tumor response in chemoresistant disease (Lengyel et al. 2020, Colombo et al. 2021).

Taken together, the increased expression of mTOR in select tumors may provide an opportunity for patient stratification or maintenance strategy. Chemotherapy mainly functions through the induction of DNA damage. DNA damage repair deficiencies are evident in nearly 50% of HGSC tumors (i.e., *BRCA1/2* mutations, Konstantinopoulos et al. 2015), specifically the reduction or loss of homologous recombination (HR) DNA repair. Consequently, chemotherapy is often a more effective treatment to target tumor cells that are deficient in DNA damage repair mechanisms. In contrast, tumors with elevated DNA damage response and HR activity are often less chemoresponsive, thereby shortening PFS. *RAD51* is a recombinase that directly contributes to HR repair via binding to singlestranded DNA and promoting strand invasion. *RAD51* is considered imperative for functional HR repair. The ability of tumor cells to form *RAD51* foci at sites of DNA damage is an effective readout of intact DNA damage repair (Compadre et al. 2023). In our findings, higher *RAD51* log fold change following chemotherapy is correlated to worse PFS (Fig S16, S17) and is one of the most important features predicting PFS (Fig. 5). Therefore, tumors that increase *RAD51* expression following chemotherapy may have enhanced DNA damage repair activity, and thus higher chemoresistance.

In our systematic analysis, we compared results on raw NanoString gene expression data vs. NanoString data that was pre-processed following standard normalization protocols. We generally saw that normalization reduced the dynamic range of pre- and post- NACT gene expression levels, thus drawing outliers closer to the rest of the data distribution (Fig 1). We found that biomarker identification via univariate Pearson correlation results between log-fold change features and PFS was partially sensitive to normalization (Fig 2). Considering both the smaller N=6 cohort and the N=83 multi-cohort, some genes showed significant Pearson correlation results regardless of normalization, but other results were sensitive to this pre-processing choice, possibly due to the influence of outliers. However, normalization of the dataset did not appear to have a strong impact on predictive model performance in either the single or multi-cohort dataset (Fig 3). Predictive performance results from regularized machine learning models might be less sensitive to outliers. These results suggest that normalization steps should always be reported in studies analyzing NanoString data and that the sensitivity of results to normalization should be tested.

Our study has some limitations. HGSC has a diversity of presentations, and our N=83 data set, while larger than most past studies, may yet still not fully represent this diversity. Where the tumor is sampled in the collection of pre- and post- NACT data is not always recorded, so spatial heterogeneity across the matched pre- and post-samples in our dataset may be an important factor. Un-measured clinical variables (e.g., menopause status, oral contraceptive use) could be associated with disease progression, but are not available in our dataset. We combined datasets across multiple sources, and hence the PFS endpoint may have been defined slightly differently in each database. Finally, for machine learning analyses, normalization should happen only within the training dataset but, in this case, data normalization steps were applied manually ahead of machine learning analysis on the entire dataset using the nSolver software, following typical procedures for working with NanoString gene expression data. This technical limitation led to a small amount of data leakage from the training to test sets in the evaluation of predictive performance, so the out-of-sample predictive performance estimates for the normalized dataset could be slightly overestimated.

This work demonstrates the value of using machine learning analyses alongside traditional univariate statistical analyses for identifying biomarkers in HGSC from high-dimensional biological data including NanoString (Clauset et al. 2021). Future work should investigate larger datasets of pre- and post- NACT matched samples by also investigating RNA-seq datasets. There are several studies that include paired pre- and post-NACT samples analyzed with RNA-seq (Adzibolosu et al. 2023, Zhang et al. 2022, Lahtinen et al. 2023, Javellana et al. 2022). Integrating NanoString and RNA-seq together for predictive modeling has not yet been thoroughly assessed, although previous literature suggests that assay-specific biases likely exist that are outside the scope of this study (Zhang et al. 2020, Rezapour et al. 2024, Rezapour et al. 2025).

Given the high heterogeneity we observed in our analysis, creating larger datasets by combining NanoString and RNA-seq datasets of paired pre- and post- NACT samples could provide further insight. Previous work has also identified subtypes of HGSC presentation (Zhang et al. 2016), and we could investigate whether there is a similar clustering of patients based on change in gene expression, cell type, and signaling features pre- and post- NACT. Finally, Jordan et al. 2020 also investigated the regulatory network between important genes, and future work could further investigate this network amongst the new set of genes identified in this work.

## Materials & Methods

### Sample preparation

For this analysis, 29 previously unpublished matched pre- and post- NACT samples (58 samples in total) were collected from patients at CU Anschutz. Collecting this data was approved under the University of Colorado’s Institutional Review Board Protocol COMIRB#18-0119. The samples were identified via the electronic medical record and the archived formalin-fixed tissues were retrieved from Pathology. Tissue slides were reviewed for representative tumor block sections by a board-certified pathologist with subspecialty training in gynecologic pathology (RJW). FFPE tissue blocks were sectioned into two 20-μm tissue-containing paraffin scrolls. RNA was extracted using the High Pure FFPET RNA Isolation Kit (Roche). RNA quantity and quality were assessed using an RNA ScreenTape on a TapeStation 4150 (Agilent Technologies). RNA concentration was determined by comparison with the RNA ladder and the percentage of RNA fragments greater than 200 bp was calculated (average 70.6%, all samples > 55%).

### NanoString PanCancer IO 360

For these 58 samples, 5μL (100-300ng) of each RNA sample was mixed with 8μL of the Master Mix (reporter codeSet and hybridization buffer). 2μL of the capture probeSet was added and the solution was mixed and spun down. It was placed in a 65°C thermocycler (Bio-Rad Laboratories Inc, Hercules, California, USA) for 16 hours. The samples were transferred to the preparation station (NanoString Technologies, Seattle, WA) with prepared nCounter Master Kit and a cartridge. The preparation station processed 12 lanes per run in approximately 2.5 to 3 hours. The cartridges were transferred to the Digital Analyzer (NanoString Technologies, Seattle, WA) for analysis. Cartridges were then scanned on the Digital Analyzer at 555 fields of view. Raw RCC files were exported for downstream analysis in nSolver Analysis Software. This data can be found using GEO accession number GSE319500.

### Constructing a dataset of pre- and post- NACT matched samples

An additional set of 54 matched pre- and post- NACT samples (108 samples in total) for which NanoString data was available were compiled from datasets associated with previously published studies. 6 of these matched samples were from Jordan et al. 2020 (PMID: 32928797), 31 were from James et al. 2022 (PMID:36131935, downloaded from GEO: GSE201600), and 17 were from Lodewijk et al. 2022 (PMID:35087175, downloaded from GEO: GSE181597). Patient age at diagnosis was also available as an additional clinical feature for 66 of the matched samples (not available for the samples from Lodewijk et al. 2022) and was considered as an additional feature in a supplemental test. In our analyses, we thus consider these matched samples to be from 4 labeled sources (PMID:32928797; CU Anschutz, unpublished; PMID:36131935; PMID:35087175).

### Pre-processing the NanoString data

Pre-processing of the NanoString data was performed using nSolver Analysis Software using the nSolver advanced analysis tool (version 4.0). For the normalized data, we included three steps across the entire dataset (see Supplemental Table 3). First, background thresholding was done according to the threshold Count (n=10).

Positive control normalization was then done using the geometric mean of POS_A-F with lanes flagged if normalization factors were outside the 0.3-3 range. Finally, reference/housekeeping normalization was done using the geometric mean with lanes flagged if normalization factors were outside the 0.1-10 range (genes: *DNAJC14*, *SDHA*, *SF3A1*, *NRDE2*, *STK11IP*, *ERCC3*, *G6PD*, *POLR2A*, *TBC1D10B*, *TBP*, *OAZ1*, *TLK2*, *PUM1*, *TMUB2*, *TFRC*, *MRPL19*, *PSMC4*, *GUSB*, *ABCF1*, *UBB*). For the non-normalized data, these normalization steps were not applied. The RLF file from nCounter PanCancer IO 360 panel was used with platform and annotation information available on GEO using GEO accession GPL27956. After removing housekeeping genes, pre- and post-measurements for the remaining 750 genes were used as features in further analyses.

The nSolver analysis tool was also used to produce 13 cell type features (Total TILs, B cells vs TILs, Cytotoxic cells vs TILs, DC vs TILs, Exhausted CD8 vs TILs, Macrophages vs TILs, Mast cells vs TILs, Neutrophils vs TILs, NK cells vs TILs, T cells vs TILs, CD8 T cells vs TILs, CD8 vs Exhausted CD8, CD4 vs T cells) and 25 signaling features (Angiogenesis, Antigen Presentation, Apoptosis, Autophagy, Cell Proliferation, Costimulatory Signaling, Cytokine and Chemokine Signaling, Cytotoxicity, DNA Damage Repair, Epigenetic Regulation, Hedgehog Signaling, Hypoxia, Immune Cell Adhesion and Migration, Interferon Signaling, JAK-STAT Signaling, Lymphoid Compartment, MAPK, Matrix Remodeling and Metastasis, Metabolic Stress, Myeloid Compartment, NF-kappaB Signaling, Notch Signaling, PI3K-Akt, TGF-beta Signaling, Wnt Signaling) for each sample on both the normalized and non-normalized datasets. All 38 of these features were used for further analysis.

### Feature engineering

The 750 gene expression and 38 cell type / signaling features for pre- and post-NACT samples were transformed into log-fold change features for each of the 83 matched samples. For the gene features, these were calculated as log2(post/pre) for the matched values pre- and post- NACT. For the cell type / signaling log-fold change features, these were calculated as post-pre for the matched values, as the features produced by the nSolver software were already log transformed. This resulted in 788 features considered for each of the matched samples.

### Outcome engineering

Our primary outcome of interest in this study is PFS, which is defined as the number of months from upfront treatment completion to first disease recurrence. This value was recorded for each of the matched samples. For those censored patients with no observed recurrence, PFS was defined as time to last patient contact. For the statistical analyses, PFS was considered a numerical value, with censoring indicators also considered for survival analysis. For our primary predictive analysis, we transformed PFS into a binary outcome variable by splitting the dataset at the median PFS of all patients (10 months, including censored patients). This binary outcome is defined as 1 for patients with a PFS value greater than the median and 0 for patients with a PFS value less than or equal to the median. Three patients in the dataset were censored before this median value, and thus have unknown binarized PFS values.

These three patients were dropped from this predictive analysis, leaving 40 patients in each PFS binary category. For additional predictive modeling performed as a sensitivity test, we consider the constructed outcome of chemotherapy status, which is defined clinically based on 3 categories (chemo-refractory=PFS less than 6 months, N=18 in our dataset; chemo-resistant=PFS greater than or equal to 6 months and less than 12 months, N=24 in our dataset; chemo-sensitive=PFS greater than or equal to 12 months; N=38 in our dataset; undefined for 3 patients censored before 12 months) (Davis et al. 2014). We consider PFS a numerical value with censoring for sensitivity tests using predictive survival analysis. Finally, 4 additional binary outcomes were also constructed for performing sensitivity tests for binary prediction within each source by splitting at their respective PFS median values. As a sensitivity test to explore whether the features were distinguishable by data source, we also performed tests predicting source from features.

### Differential expression and statistical analyses

We ran differential expression and statistical analyses to replicate the same analyses performed in prior studies on this larger combined dataset. Differential expression, Pearson correlation, Kaplan-Meier, and Cox regression analyses were performed in R (version 4.4.3). Cox regression and Kaplan-Meier analyses used the survival package (version 3.8.3, Therneau 2025).

### Machine learning analyses

Machine learning predictive analyses for binary, chemotherapy status, and source outcomes were run in Python (version 3.12.18) using the sckit-learn library (version 1.6.1). Features were separately scaled in the training and test sets using StandardScaler. Logistic regression was explored over three penalties (L1, L2, and ElasticNet) using the saga solver. The L1 ratio was set to 0.5 for the ElasticNet penalty. The random forest used default hyperparameters. Predictive performance on the test set was evaluated via the ROC-AUC, Brier score, accuracy, precision, recall, and F1 score metrics. ROC-AUC on the training dataset was also evaluated. Model performance was averaged over 10 iterations of leave-one-out cross validation (LOOCV). For each of these 10 iterations of LOOCV, an independent randomized version of the data was also produced for accompanying evaluation of performance on data with randomly permuted feature columns.

For multi-class classification of chemotherapy status and data source, performance metrics based on predicted class (precision, recall, F1 score) were combined over multi-class outcomes using the weighted average strategy. For sensitivity tests exploring logistic regression hyperparameters we tuned over 20 options for the regularization hyperparameter using 5-fold cross validation optimizing ROC-AUC, with performance results averaged over 5-fold cross validation. For sensitivity tests exploring random forest hyperparameters, we tested 64 randomly selected combinations of default and extreme values for max_features, min_sample_split, n_estimators, max_depth, and min_samples_leaf (see Fig. S7). For sensitivity tests with Age included as an additional feature, simple mean imputation was used to impute missing Age values for Lodewijk et al. 2022. For sensitivity tests using machine learning for survival analysis, Cox proportional hazards models were run using CoxnetSurvivalAnalysis from the scikit-survival library (version 0.24.1, Pölsterl 2020) with a L1 ratio of 0.9. Random survival forests were run using the randomForestSRC package (version 3.5.1, Ishwaran and Kogalur 2007, Ishwaran et al. 2008, Ishwaran and Kogalur 2025) in R with the hyperparameters nodesize and mtry tuned to optimal values based on the out-of-sample error, ntree=100, and using the log-rank spit rule to split on the largest difference between survival distributions of two groups. For survival machine learning models, we evaluated performance based on Harrell’s C index.

### Feature importance investigations

Feature importance values were extracted from the trained random forest models based on Gini importance (Nembrini et al. 2018) and from the logistic regression models based on the absolute value of model coefficients and averaged across folds and iterations of model training. The results across normalized and non-normalized datasets for the 4 models encompassing the random forest and 3 versions of logistic regression based on the L1, L2, and Elastic penalties produced 8 ranked lists of feature importances. We combined these 8 lists into a final aggregated list of top features using the Borda method for rank aggregation (Sarkar et al. 2014) and considered the top 20 from this aggregated list as the top features identified from the predictive models.

As a sensitivity test for these aggregated feature importance results, we also considered the top 20 features from multivariable Cox proportional hazards models for both normalized and non-normalized data.

### Principal component analysis

Principal component analysis (PCA) was performed in Python (version 3.12.18) using the sckit-learn library (version 1.6.1).

### Ethics statement

Formalin fixed tissue blocks were acquired (COMIRB#18-0119) and a board-certified pathologist confirmed tumor tissue. This protocol was deemed exempt, as it used previously collected data, the information was not recorded in a manner that is identifiable, and the gained information did not change clinical decision making.

## Data Availability Statement

The 29 new matched samples have been deposited at GEO: GSE319500 along with the 6 matched samples from Jordan et al. 2020. 31 matched samples were downloaded from GEO: GSE201600 (James et al. 2022) and 17 were downloaded from GEO: GSE181597 (Lodewijk et al. 2022).

## Acknowledgements

We acknowledge philanthropic contributions from Kay L. Dunton Endowed Memorial Professorship in Ovarian Cancer Research. We thank Esther Rolf, Robbe D’hondt, and Celine Vens for helpful conversations. We acknowledge funding from the Ovarian Cancer Research Alliance Collaborative Award (Clauset A and Bitler B), NIH/NCI R37 (Bitler B), R01 (Bitler B and Sikora M), R50CA293845 (Jordan K), DOD OCRP OC240039 (Bitler B), the University of Colorado Cancer Center Support Grant (P30CA046934). We also thank the Human Immune Monitoring Shared Resource (RRID:SCR_021985).

## Author contributions

Lucy B. Van Kleunen: Conceptualization, data curation, software, formal analysis, validation, investigation, visualization, methodology, writing–original draft, writing–review and editing.

Grace Bowman: Data curation, software, formal analysis, validation, investigation, visualization, methodology, writing–original draft, writing–review and editing.

Sarah Elizabeth Stockman: Data curation, software, formal analysis, validation, investigation, visualization, methodology, writing–original draft, writing–review and editing.

Logan Barrios: Data curation, software, formal analysis, validation, investigation, visualization, methodology, writing–original draft, writing–review and editing.

Hope A. Townsend: Data curation, methodology, writing–original draft, writing–review and editing.

Kimberly R. Jordan: Resources, data curation, funding acquisition, validation, investigation, methodology, writing–review and editing.

Rebecca J. Wolsky: Data curation, validation, writing–review and editing.

Kian Behbakht: Conceptualization, funding acquisition, writing–review and editing. Matthew J. Sikora: Conceptualization, writing–review and editing.

Jennifer K. Richer: Writing–review and editing.

Benjamin G. Bitler: Conceptualization, resources, supervision, funding acquisition, validation, investigation, visualization, methodology, writing–original draft, writing–review and editing, project administration,

Junxiao Hu: Validation, investigation, methodology, writing - review and editing.

Aaron Clauset: Conceptualization, resources, supervision, funding acquisition, validation, investigation, visualization, methodology, writing - original draft, writing–review and editing, project administration.

