## Supporting Information for "On the predictability of progression-free survival in ovarian cancer from NanoString gene expression data"

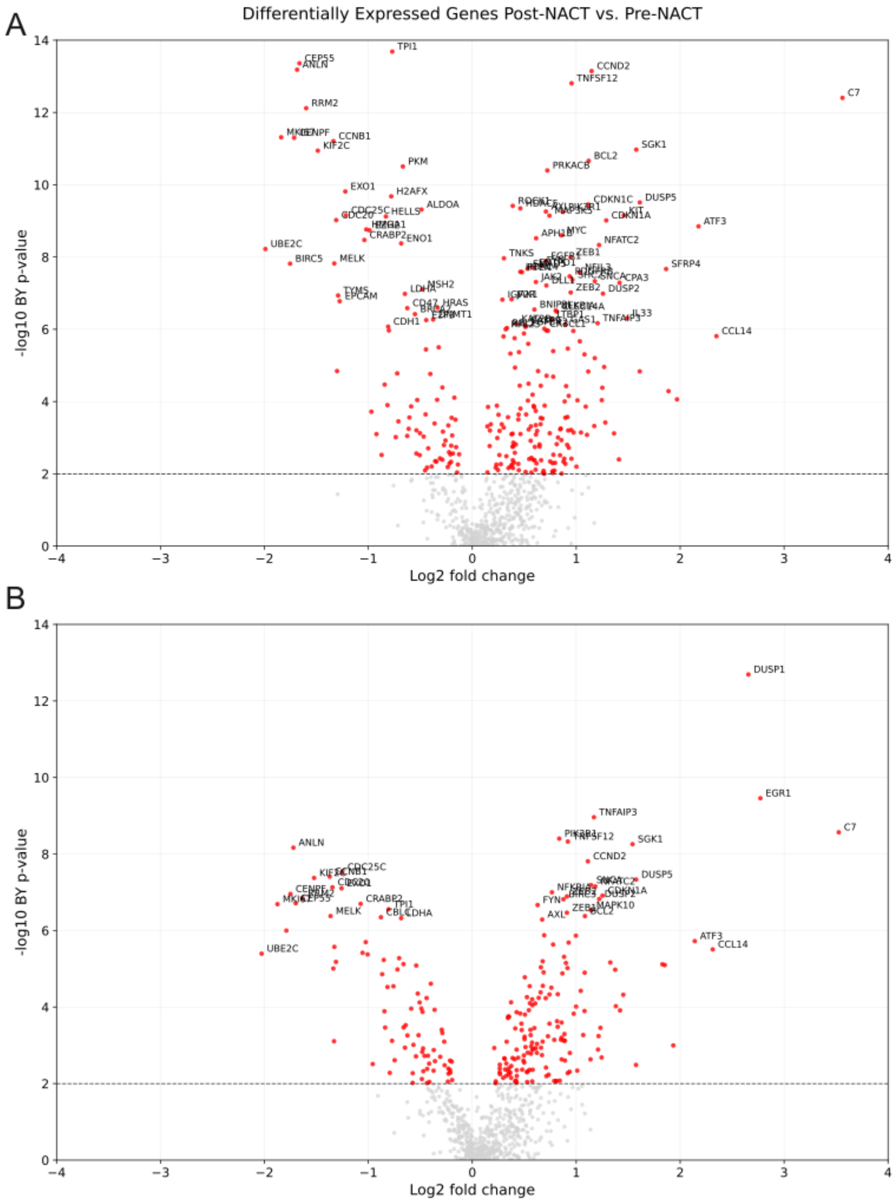


**SI Figure 1.** Volcano plots showing differential expression pre- and post- NACT for (A) normalized and (B) non-normalized versions of the dataset. P-values are based on paired two-sided t-tests, with results highlighted in red for which the p-value was less than 0.01. Select genes with either high p-values (-log10(p-value) > 6) or high log2 fold change values (log2 fold change > 2) are labeled.

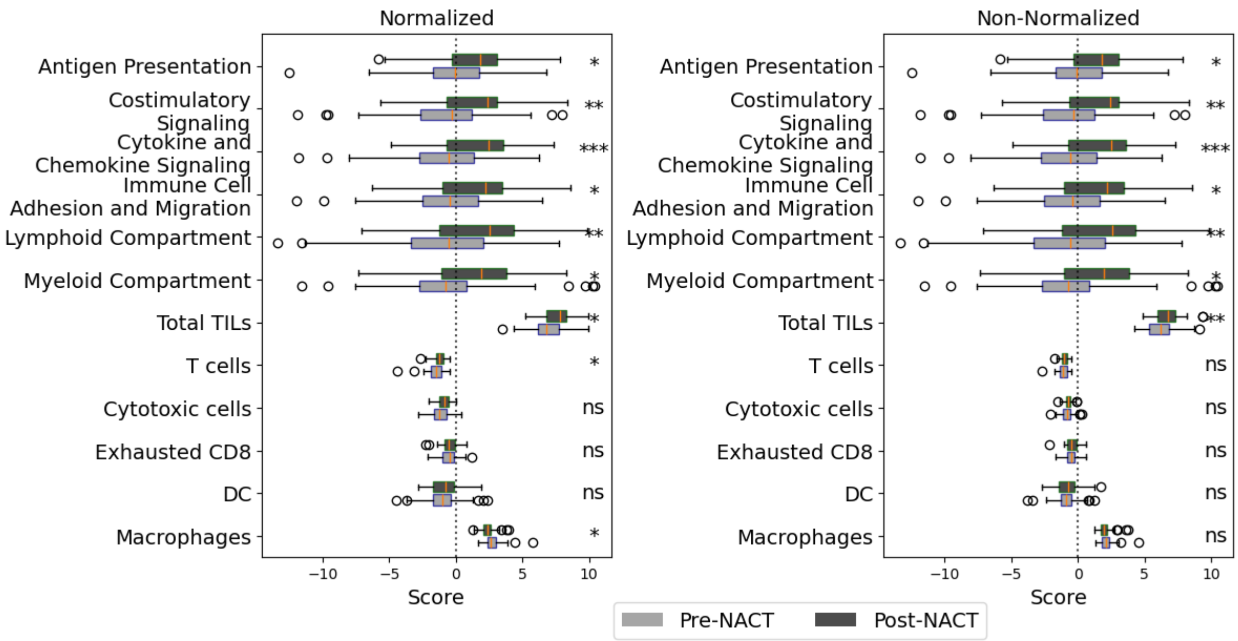


**SI Figure 2.** Differences pre- and post- NACT for the 12 cell-type and signaling scores found to increase post-NACT in Lodewijk et al. 2022 for (A) normalized and (B) non-normalized data for N=42 patients with worse prognosis representing an inflammatory signature (PFS < 12 months). Significance is determined by paired t-tests (non-adjusted, *** p< 0.001, ** p<0.01, *p<0.05, ns=not significant).

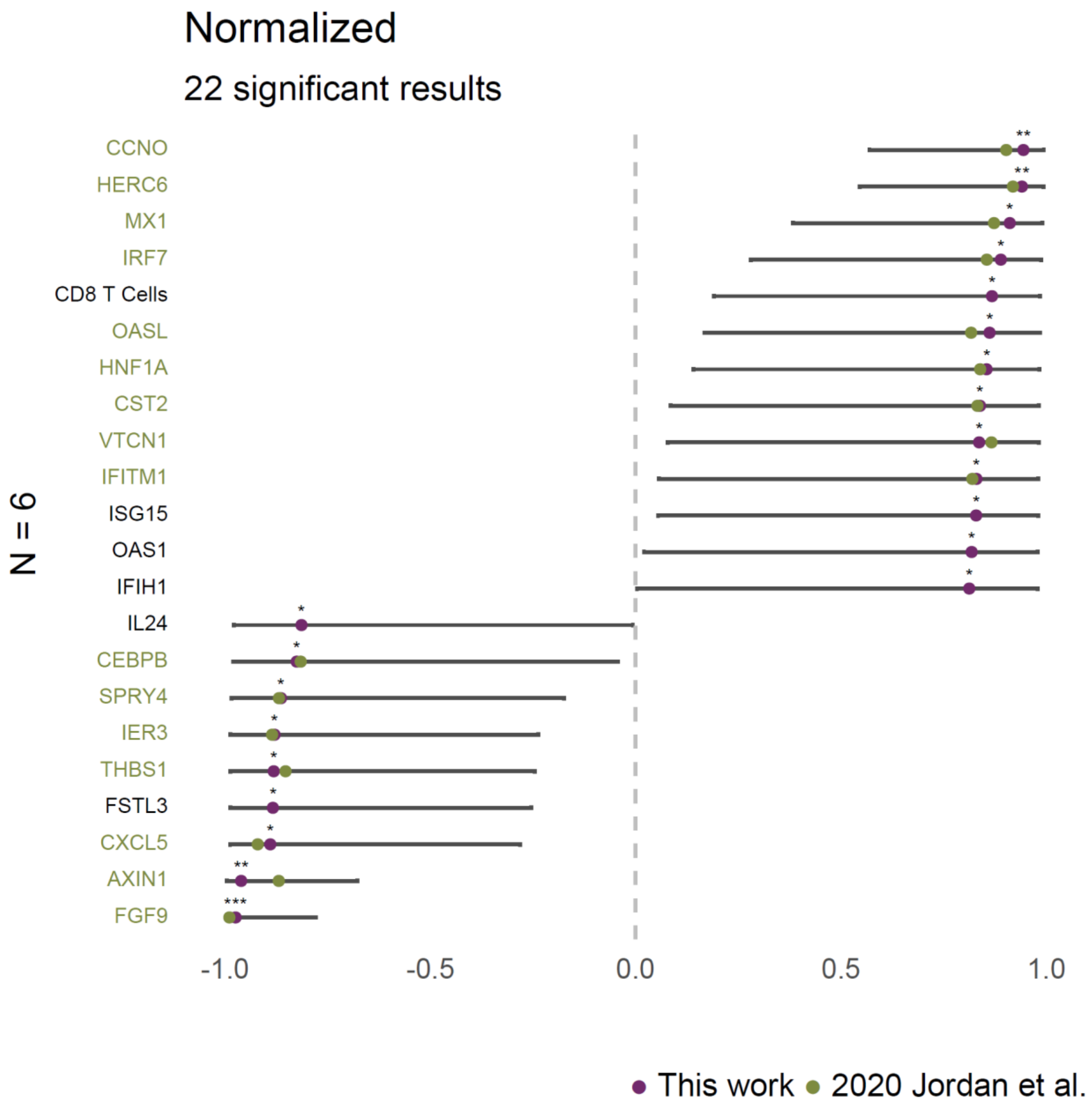


**SI Figure 3.** Significant results from Pearson correlation analysis between 788 gene, cell, and signaling log-fold change features and PFS performed for the N=6 samples from Jordan et al. 2020 on the version of the dataset normalized across the N=83 samples. Asterisks indicate significance levels (non-adjusted, **** p< 0.0001, *** p< 0.001, ** p<0.01, *p<0.05). Green dots indicate estimates for features in the original Jordan et al. 2020 analysis and feature labels are colored green if they were significant in the Jordan et al. 2020 results (across 750 gene features). 16/18 results replicated, while 2 were no longer significant under this alternative normalization: *ITGAV*, *CASP3*.

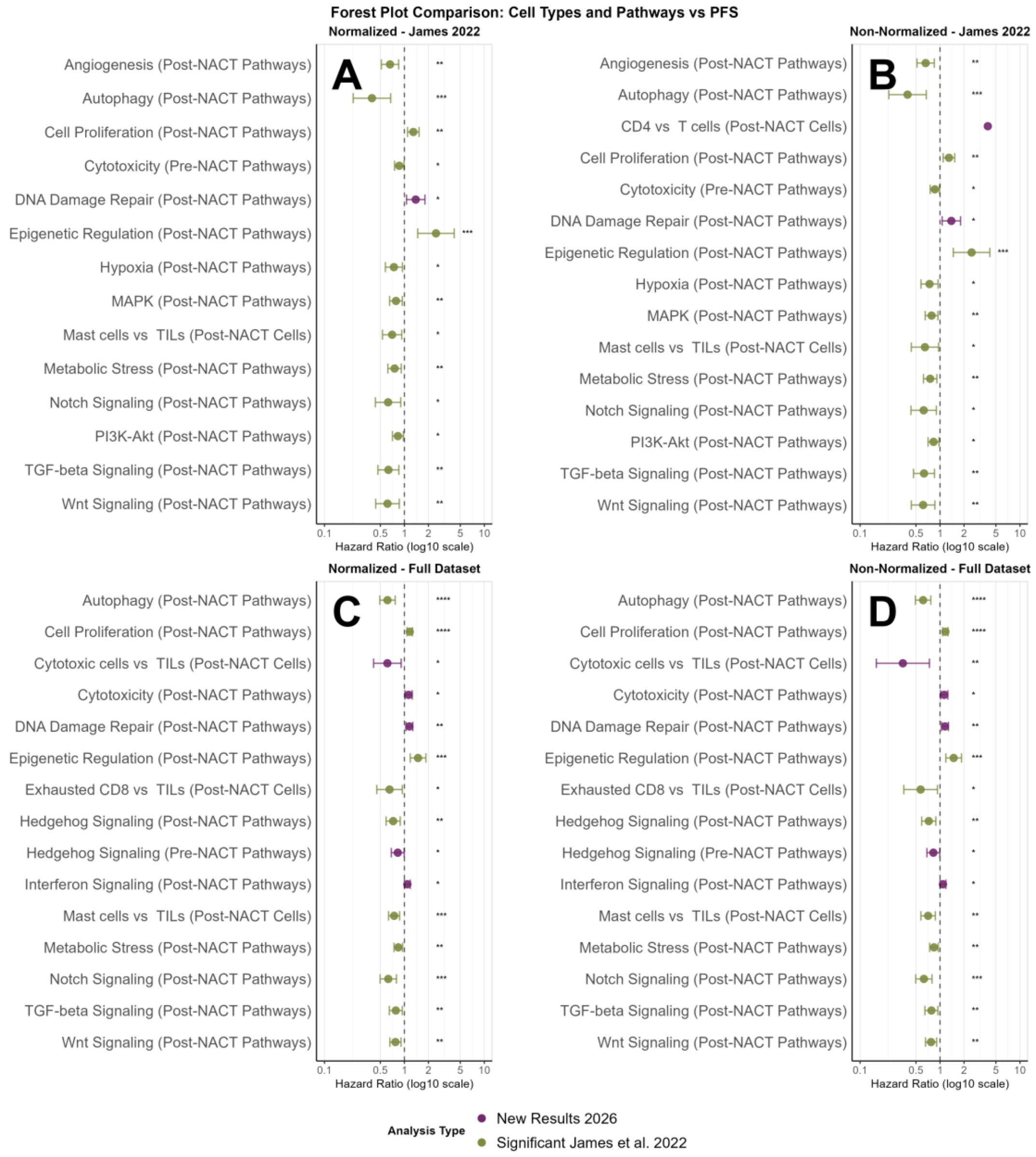


**SI Figure 4.** Univariate Cox regression results performed for pre- and post- cell type and signaling features. (A) For the N=31 samples from James et al. 2022, normalized (normalization in this case was performed on the N=83 dataset, thus not completely replicating the data pre-processing from the original paper). (B) For the N=31 samples from James et al. 2022, non-normalized. (C) For the N=83 dataset, normalized. (D) For the N=83 dataset, non-normalized. Hazard ratio estimates are shown on a log 10 scale, with a vertical line at 1, indicating neither a positive or negative impact on survival. Estimates are colored green if they were also found to be significant in James et al. 2022. Stars indicate significance levels (non-adjusted, **** p< 0.0001, *** p< 0.001, ** p<0.01, *p<0.05). Significant features are listed in alphabetical order. All results which were previously significant in James et al. 2022 were found to have the same direction of effect in these analyses.

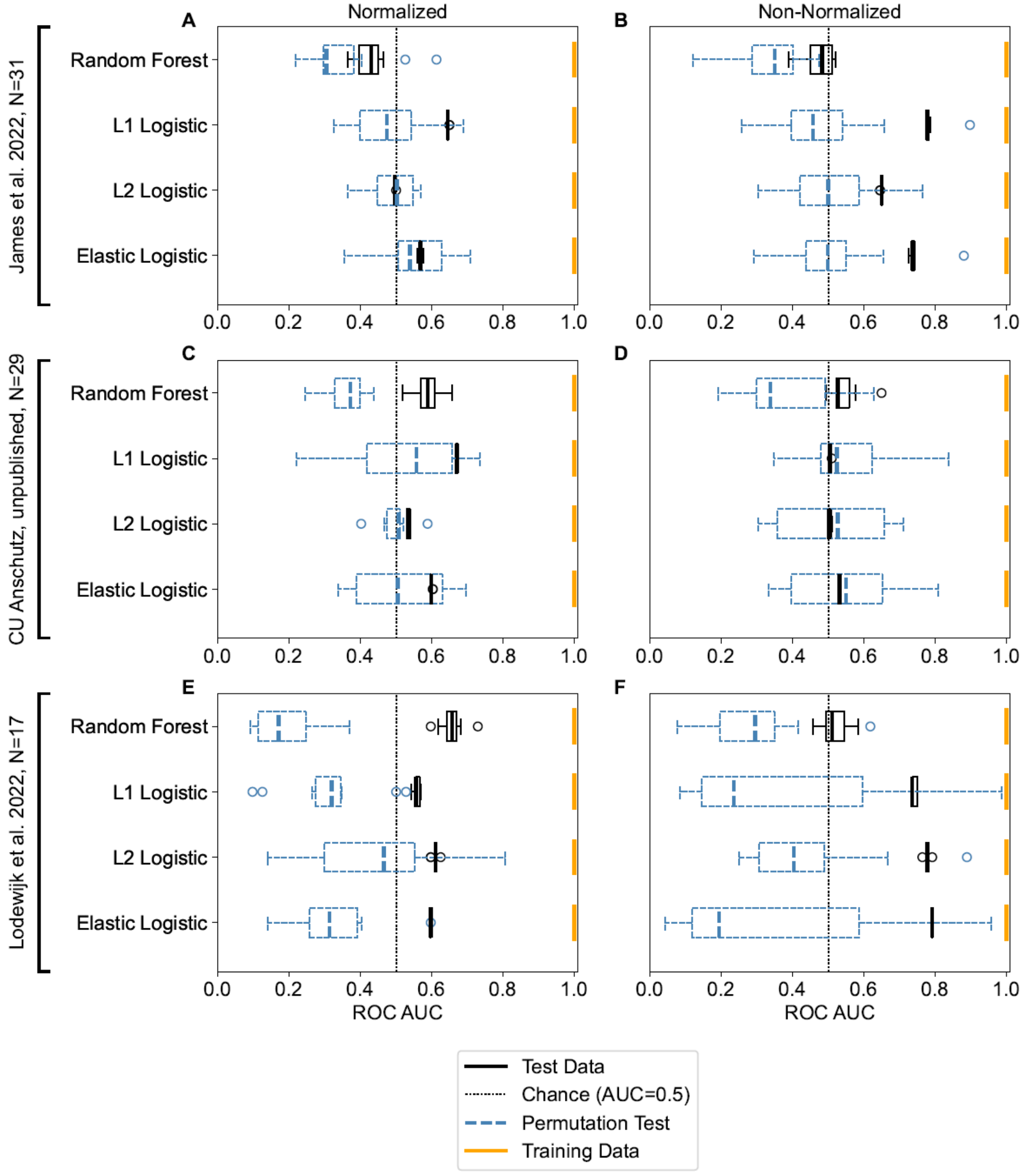


**SI Figure 5.** Predictive performance results for binary classification of PFS based on the ROC-AUC metric where PFS is split at each respective data source median and prediction is performed entirely in each data source, reported over 10 iterations of leave-one-out cross-validation. (A) For the N=31 samples from James et al. 2022, normalized, (B) for the N=31 samples from James et al. 2022, non-normalized, (C) for the N=29 unpublished samples from CU Anschutz, normalized (D) for the N=29 unpublished samples from CU Anschutz, non-normalized, (E) for the N=17 samples from Lodewijk et al. 2022, normalized, (F) for the N=17 samples from Lodewijk et al. 2022, non-normalized. The x axis of the plot ranges from 0 to 1 based on the range of possible ROC-AUC values, and a vertical dashed line is shown at ROC-AUC=0.5, which represents a baseline value above which predictive performance is better than a random guess. The distribution of results on the test dataset across the iterations is shown in black. The mean performance on the training dataset across iterations is shown in yellow. The distribution of results from dataset-specific feature permutation tests are shown in blue (dashed). Box plots show the median and the whiskers represent 1.5 times the interquartile range.

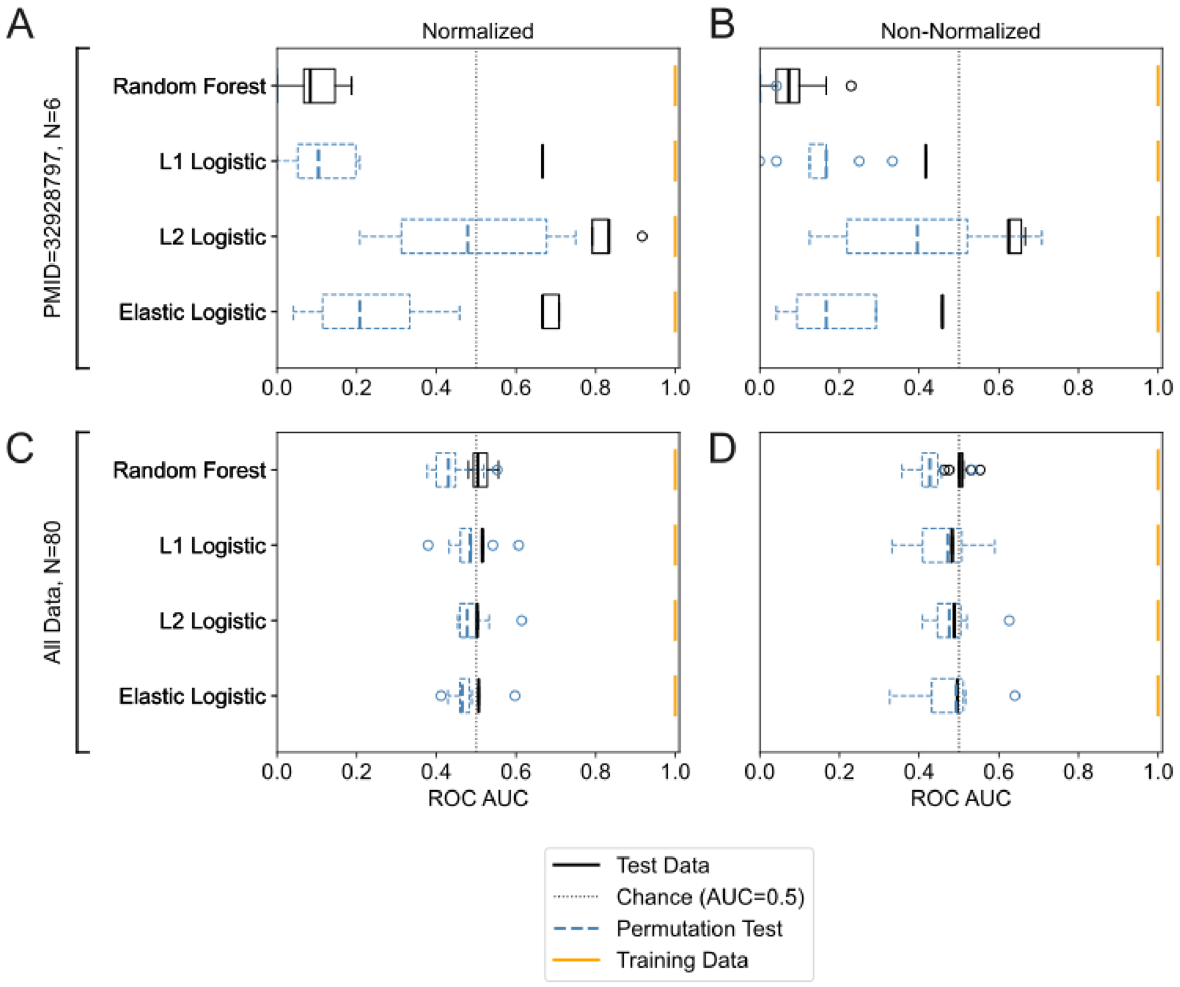


**SI Figure 6.** Multiclass prediction of chemotherapy status based on the AUC metric. (A) For the N=6 samples from Jordan et al. 2020 on the normalized dataset, (B) for the N=6 samples from Jordan et al. 2020, non-normalized, (C) for the N=80 samples, normalized, and (D) for the N=80 samples, non-normalized. Normalized versions of the data are normalized over all N=83 samples in the dataset. The x axis of the plot ranges from 0 to 1 based on the range of possible ROC-AUC values, and a vertical dashed line is shown at ROC-AUC=0.5, which represents a baseline value above which predictive performance is better than a random guess. The distribution of results on the test dataset across the iterations is shown in black. The mean performance on the training dataset across iterations is shown in yellow. The distribution of results from dataset-specific feature permutation tests are shown in blue (dashed). Box plots show the median and the whiskers represent 1.5 times the interquartile range.

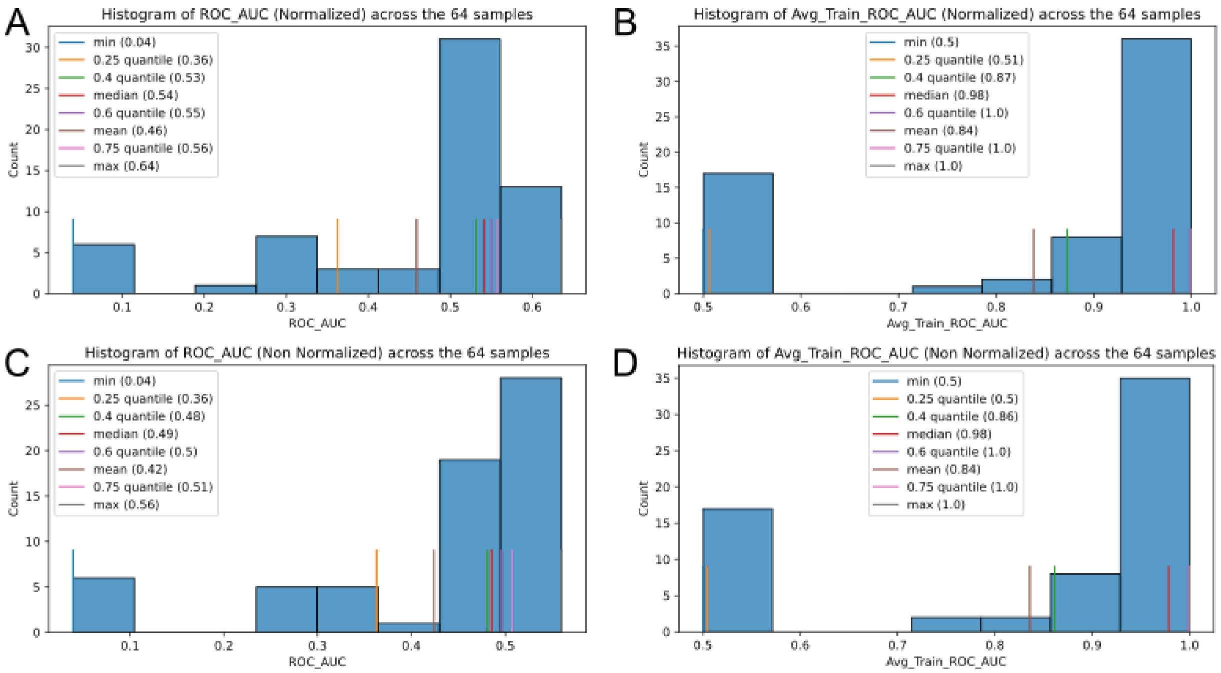


**SI Figure 7.** Results of random forest hyperparameter tests to assess whether alternative random forest hyperparameter combinations could improve predictive performance for binary classification of low vs. high PFS on the N=80 dataset based on all 788 log fold change features. We explored the space of hyperparameters to attempt to find a regime that would show improved performance. We tested the following hyperparameters over library defaults (bolded) and values we expected might influence performance based on prior literature: n_estimators=[5, 10, **100**, 2000], max_features =[0.01, **sqrt**, 0.1], min_sample_split=[**2**, 5, 10], max_depth=[1, 3, 5, **None**], and min_samples_leaf=[**1**,3,5,30]. LOOCV was evaluated over 1 iteration for each of 64 different hyperparameter combinations. The combinations were chosen by testing for each hyperparameter option once with the defaults of all other hyperparameters and generating three more combinations per hyperparameter option with the values of the other hyperparameters randomly sampled from the sets of potential values, with duplicate hyperparameter combinations removed. Results are shown for (A) test performance on the normalized data, (B) train performance on the normalized data, (C) test performance on the non-normalized data, and (D) train performance on the non-normalized data. The maximum test dataset performance we found was on the normalized data, with AUC=0.64, which was close to the best average model performance we saw in our experiments, indicating poor random forest performance was likely driven by lack of a strong signal in the data rather than not performing hyperparameter selection. We found hyperparameter combinations for which AUC < 1 on the training dataset while test performance remained low, indicating that addressing random forest overfitting did not improve performance.

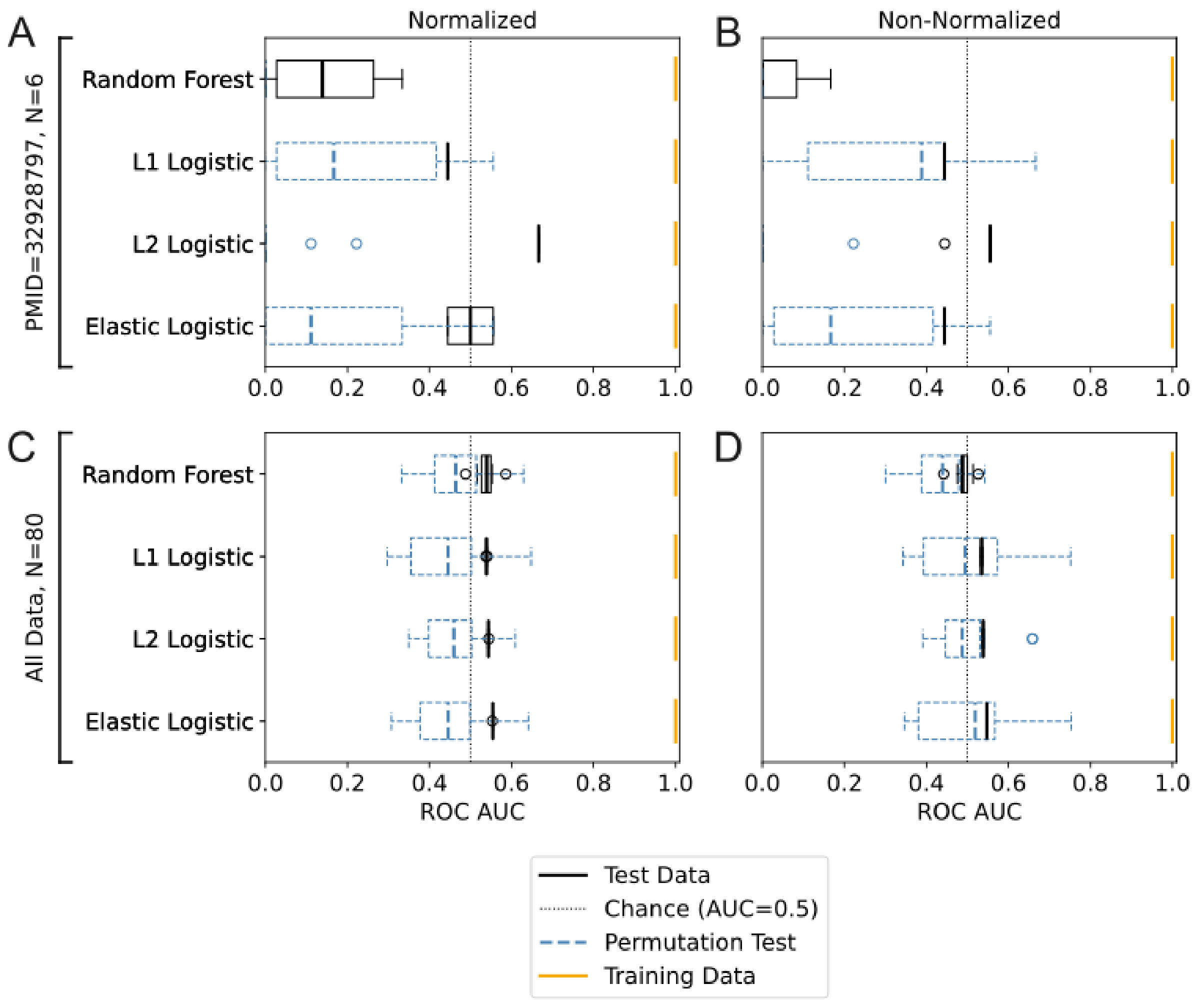


**SI Figure 8.** Predictive performance results for binary classification of PFS based on the ROC-AUC metric without feature scaling on the (A) Jordan et al 2020 dataset, normalized, (B) Jordan et al 2020 dataset, non-normalized, (C) N=80 dataset, normalized (D), N=80 dataset, non-normalized. The x axis of the plot ranges from 0 to 1 based on the range of possible ROC-AUC values, and a vertical dashed line is shown at ROC-AUC=0.5, which represents a baseline value above which predictive performance is better than a random guess. The distribution of results on the test dataset across the iterations is shown in black. The mean performance on the training dataset across iterations is shown in yellow. The distribution of results from dataset-specific feature permutation tests are shown in blue (dashed). Box plots show the median and the whiskers represent 1.5 times the interquartile range.

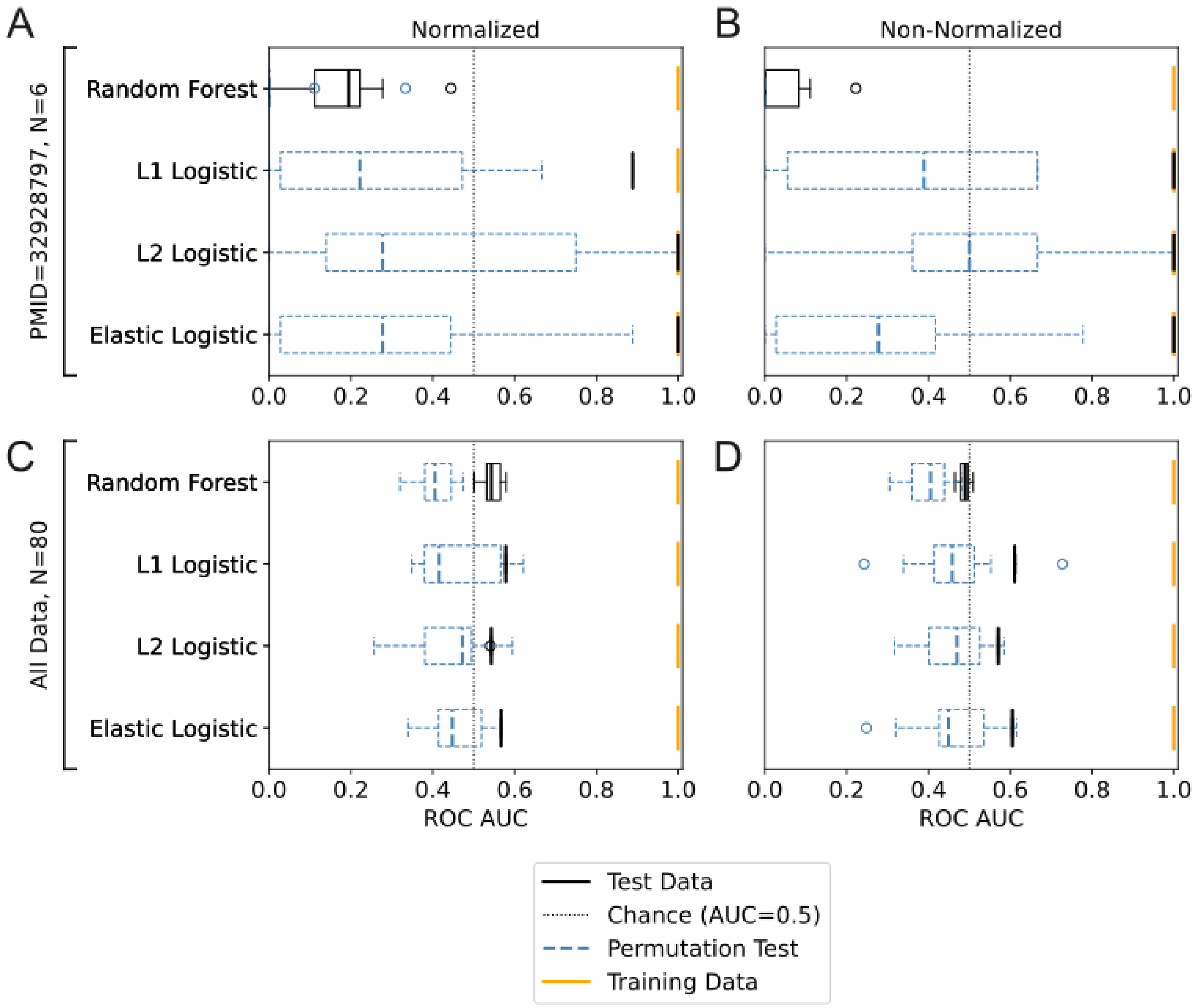


**SI Figure 9.** Predictive performance results for binary classification of PFS based on the ROC-AUC metric with Age included as an additional feature on the (A) Jordan et al 2020 dataset, normalized, (B) Jordan et al 2020 dataset, non-normalized, (C) N=80 dataset, normalized (D), N=80 dataset, non-normalized. Mean imputation was used to fill in Age for the samples for which it was not available. The x axis of the plot ranges from 0 to 1 based on the range of possible ROC-AUC values, and a vertical dashed line is shown at ROC-AUC=0.5, which represents a baseline value above which predictive performance is better than a random guess. The distribution of results on the test dataset across the iterations is shown in black. The mean performance on the training dataset across iterations is shown in yellow. The distribution of results from dataset-specific feature permutation tests are shown in blue (dashed). Box plots show the median and the whiskers represent 1.5 times the interquartile range.

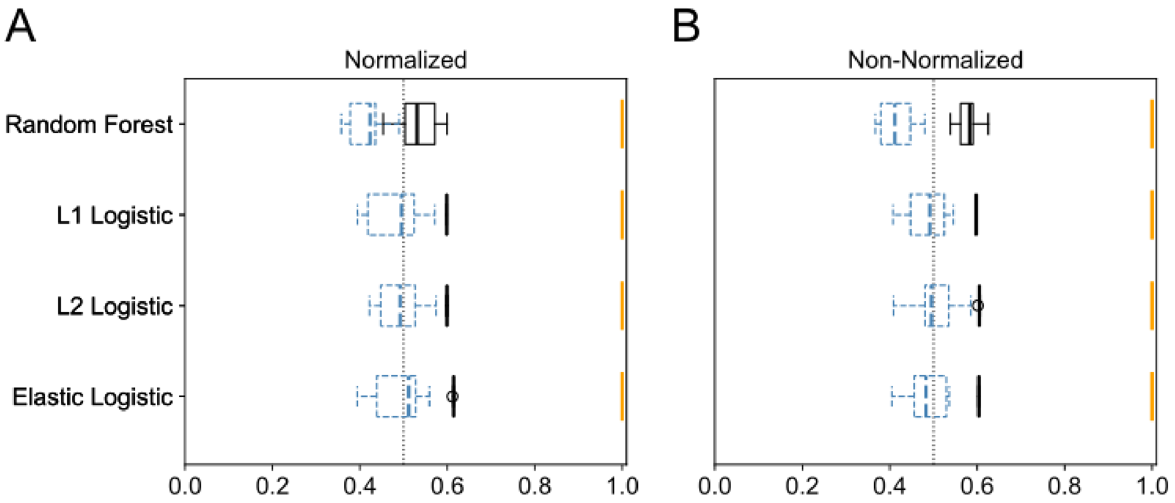


**SI Figure 10.** Predictive performance results for multiclass classification of data source based on the ROC-AUC metric across 10 iterations of leave-one-out cross-validation. (A) for the N=83 samples, normalized, and (B) for the N=83 samples, non-normalized. The x axis of the plot ranges from 0 to 1 based on the range of possible ROC-AUC values, and a vertical dashed line is shown at ROC-AUC=0.5, which represents a baseline value above which predictive performance is better than a random guess. The distribution of results on the test dataset across the iterations is shown in black. The mean performance on the training dataset across iterations is shown in yellow. The distribution of results from dataset-specific feature permutation tests are shown in blue (dashed). Box plots show the median and the whiskers represent 1.5 times the interquartile range.

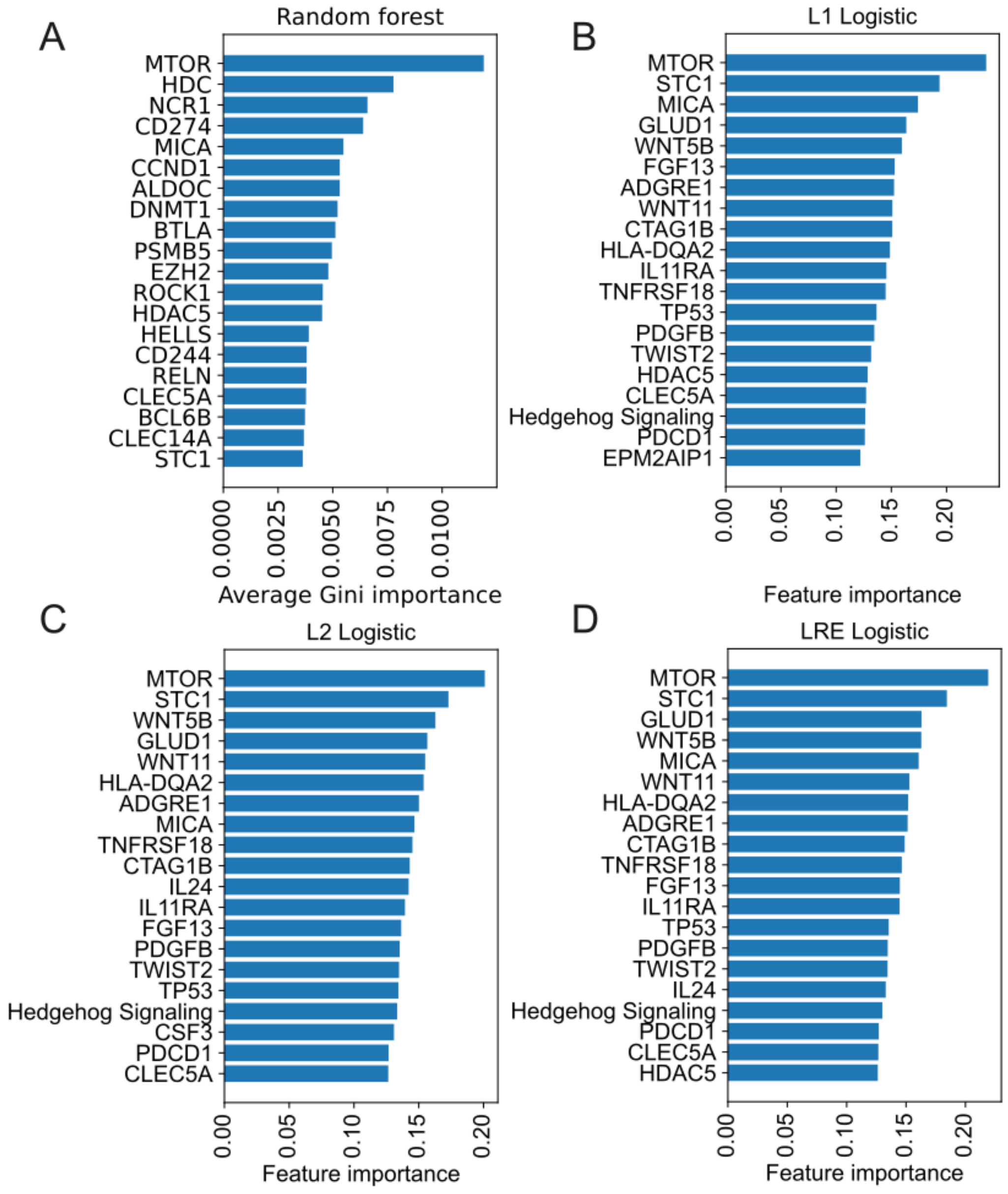


**SI Figure 11.** The 20 features with highest importance per model based on binary PFS classification on the N=80 normalized data with (A) random forest, (B) logistic regression (L1), (C) logistic regression (L2), (D) logistic regression (elastic net).

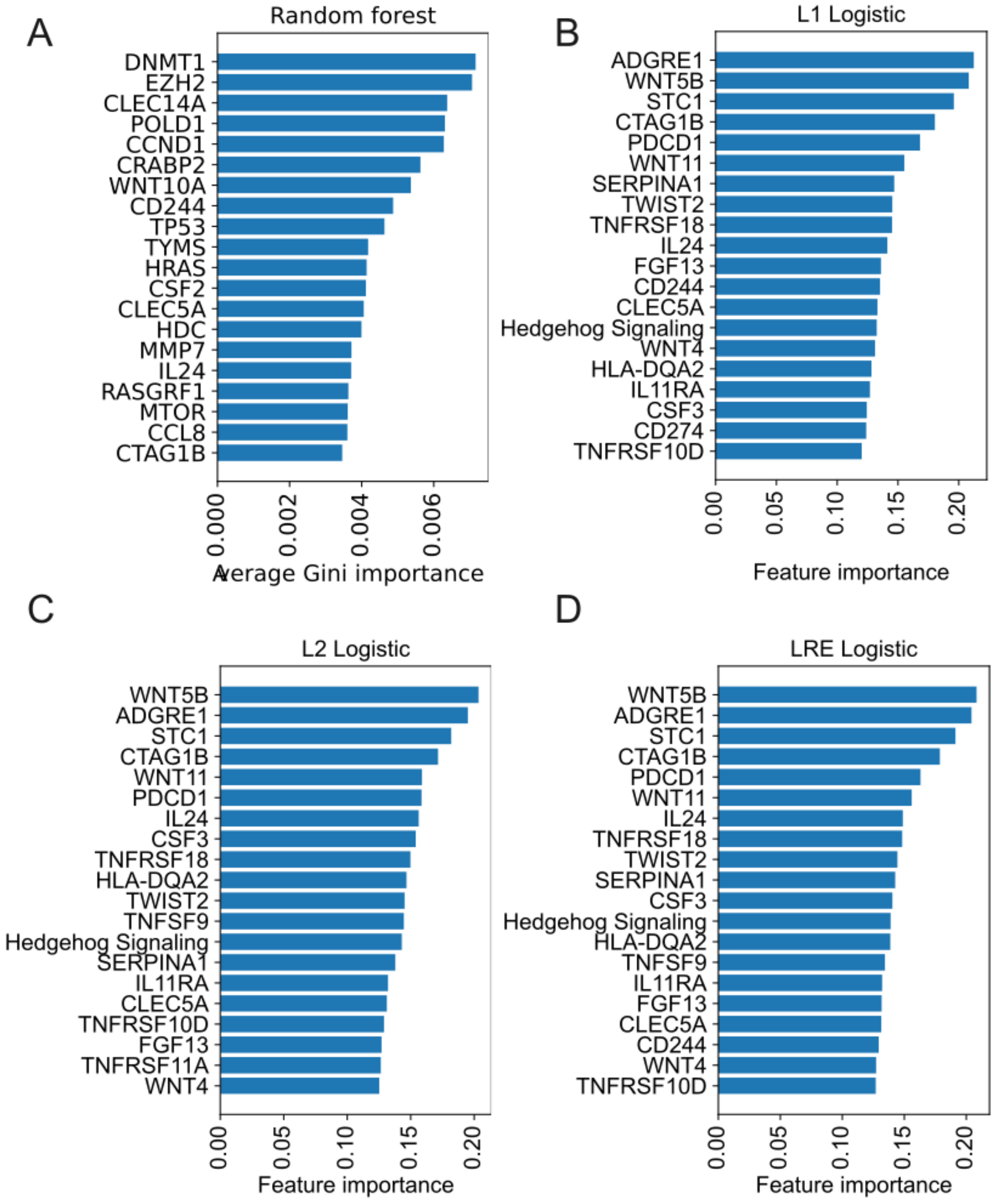


**SI Figure 12.** The 20 features with highest importance per model based on binary PFS classification on the N=80 non-normalized data with (A) random forest, (B) logistic regression (L1), (C) logistic regression (L2), (D) logistic regression (elastic net).

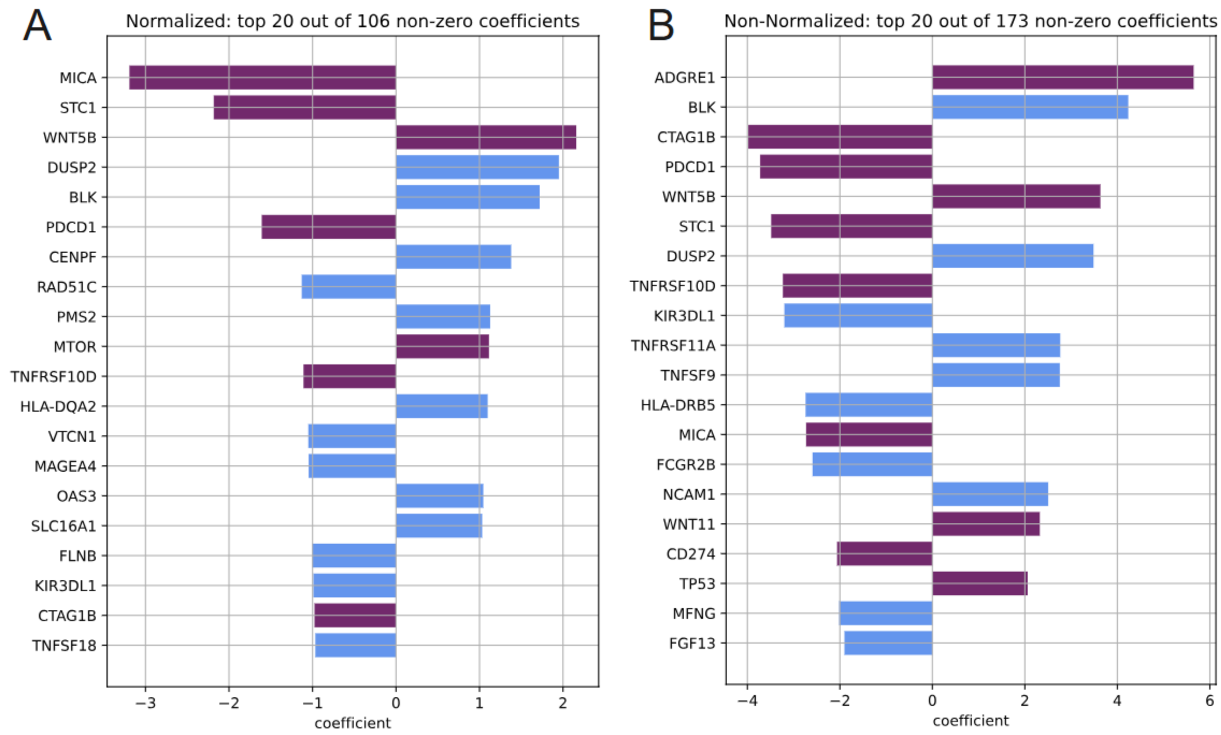


**SI Figure 13.** The 20 most important features from a regularized Cox proportional hazards model modeling PFS across the entire N=83 dataset, with features ranked based on the absolute value of coefficients. The L1 ratio was set to 0.9 and regularization strength was chosen based on 5-fold cross validation. Results are shown for (A) normalized and (B) non-normalized data. A positive coefficient indicates worse prognosis with an increase in a gene expression post-NACT, and a negative coefficient indicates better prognosis. Features are indicated with a purple bar if they are also in the set of top 20 features identified from binary prediction models reported in Figure 5 and otherwise indicated with a blue bar.

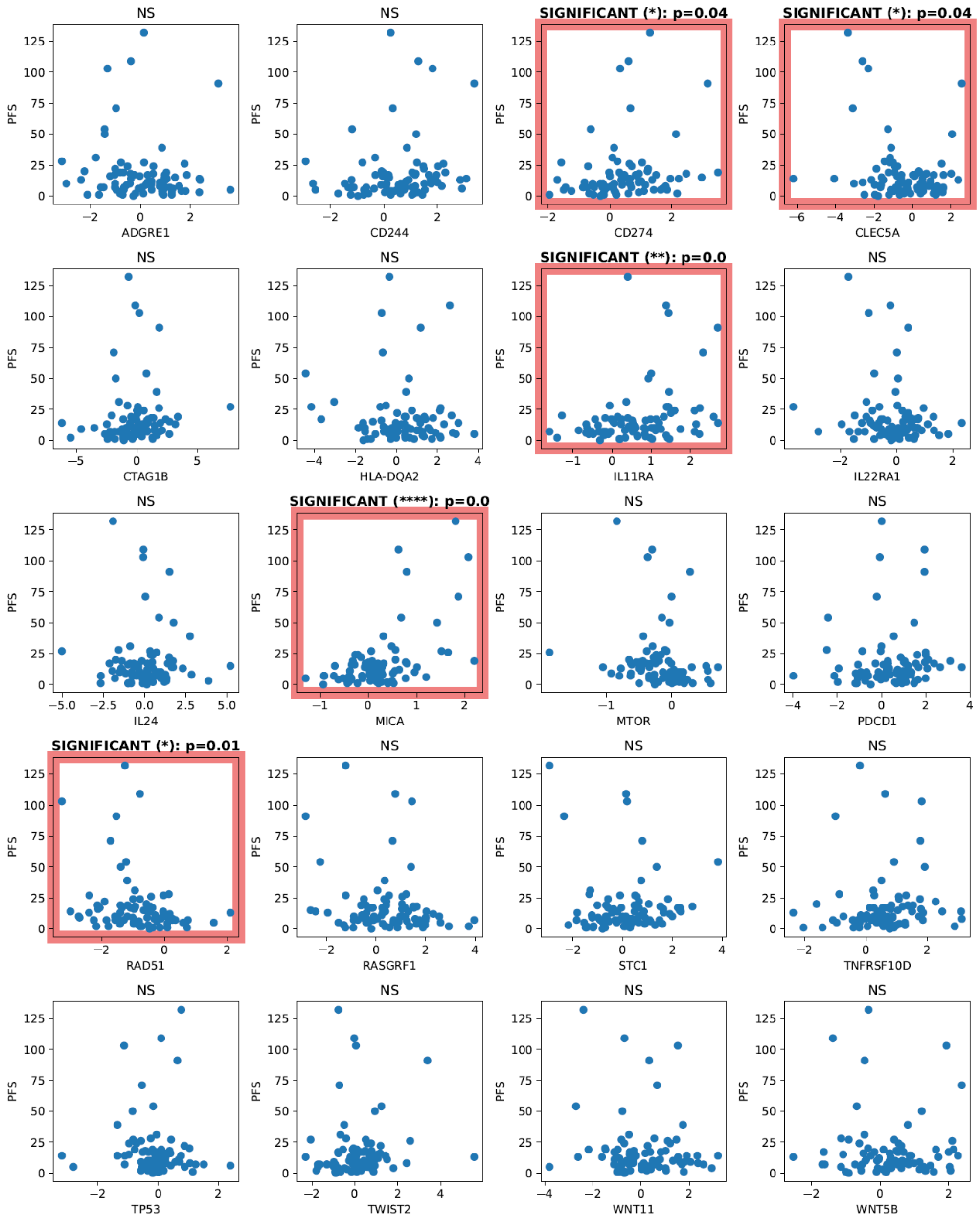


**SI Figure 14.** Scatter plots based on the full normalized dataset for the top 20 log fold change features identified from binary prediction models reported in Figure 5 vs. PFS showing significance of Pearson correlation analysis (non-adjusted, NS= not significant, **** $p < 0.0001$, *** $p < 0.001$, ** $p<0.01$, *$p<0.05$). 5 of these features had significant Pearson correlations with PFS.

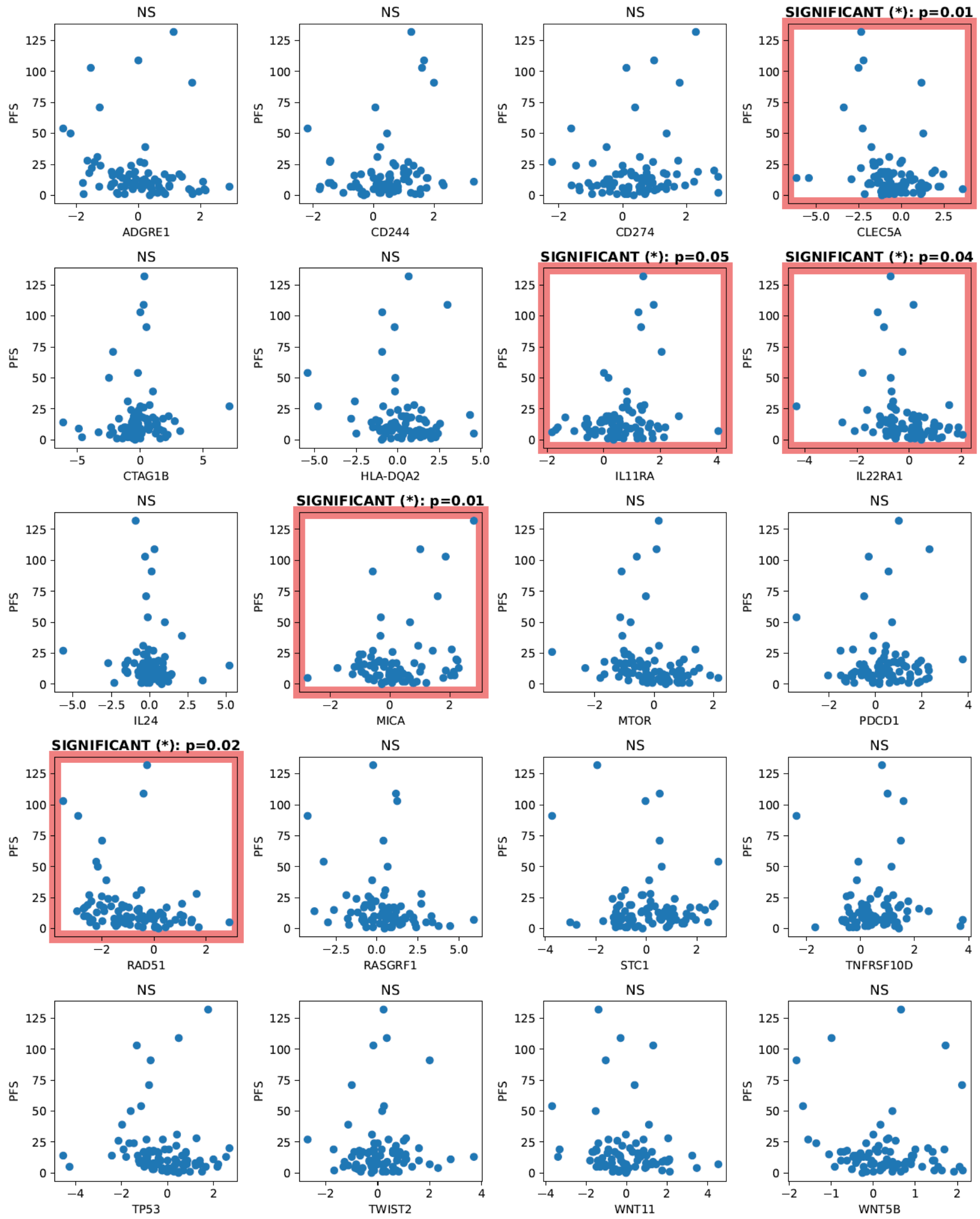


**SI Figure 15.** Scatter plots based on the full non-normalized dataset for the top 20 log fold change features identified from binary prediction models reported in Figure 5 vs. PFS showing significance of Pearson correlation analysis (non-adjusted, NS= not significant, **** p< 0.0001, *** p< 0.001, ** p<0.01, *p<0.05). 5 of these features had significant Pearson correlations with PFS.

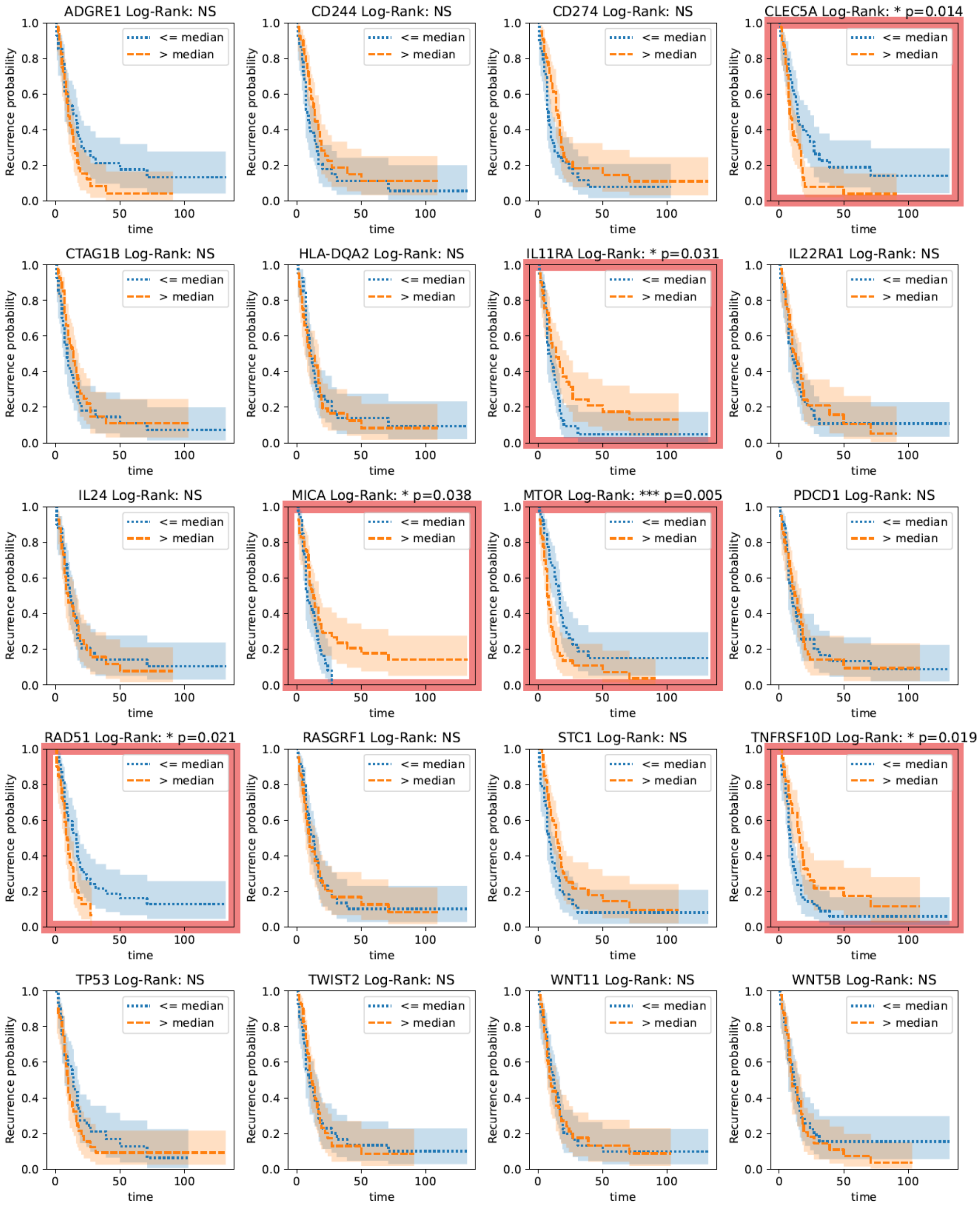


**SI Figure 16.** Kaplan-Meier curves based on the full normalized dataset for the top 20 log fold change features identified from binary prediction models reported in Figure 5. The population is split at the median feature value and survival curves are compared via the log-rank test, for which significance is shown (non-adjusted, NS= not significant, **** p< 0.0001, *** p< 0.001, ** p<0.01, *p<0.05). Six of these features had significant log-rank tests.

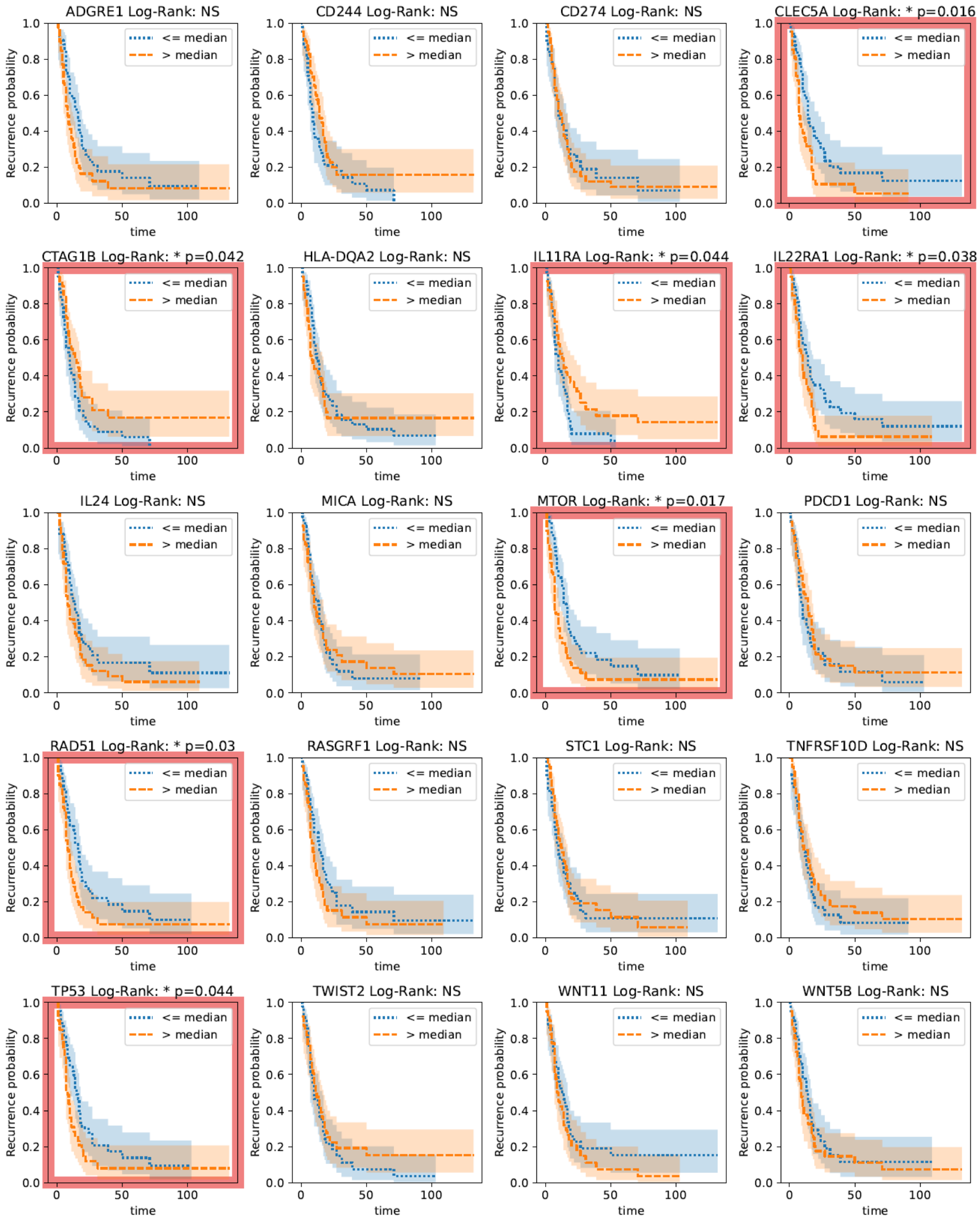


**SI Figure 17.** Kaplan-Meier curves based on the full non-normalized dataset for the top 20 log fold change features identified from binary prediction models reported in Figure 5. The population is split at the median feature value and survival curves are compared via the log-rank test, for which significance is shown (non-adjusted, NS= not significant, **** p< 0.0001, *** p< 0.001, ** p<0.01, *p<0.05). Seven of these features had significant log-rank tests.

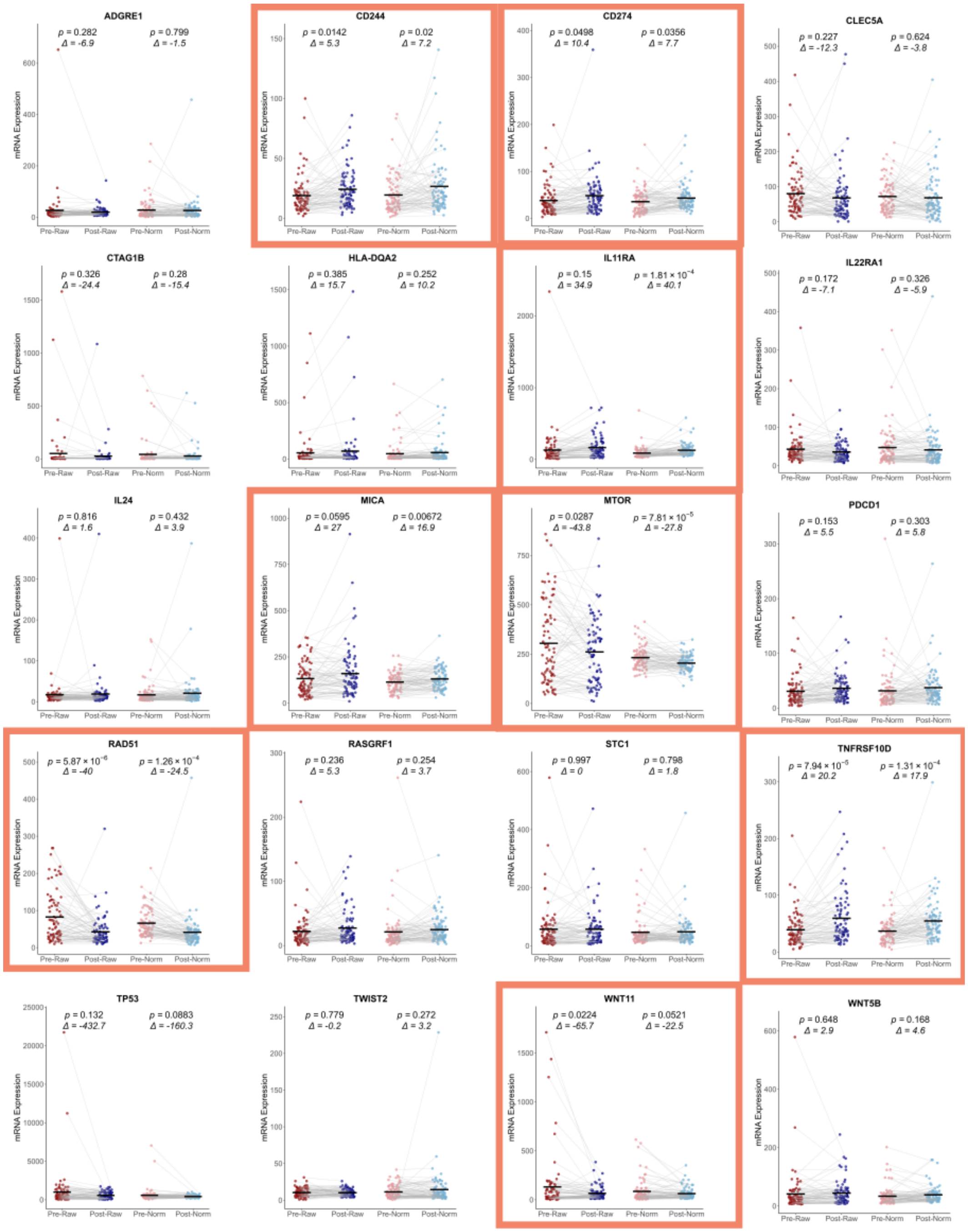


**SI Figure 18.** The global shift pre- and post- NACT for the top 20 log fold change features identified from binary prediction models reported in Figure 5. Eight of these genes which also showed a global shift significant for either raw (non-normalized) or normalized data are highlighted.

**Supplemental Tables**

- **Supplemental Table 1**: Differential expression pre- vs. post- NACT results for all 788 features on the N=83 dataset for normalized and non-normalized data.
- **Supplemental Table 2**: Normalization constants for the N=6 normalized data
- **Supplemental Table 3**: Normalization constants for the N=83 normalized data
- **Supplemental Table 4**: Pearson correlation results for 788 log-fold change features and PFS on the N=83 dataset for normalized and non-normalized data.
- **Supplemental Table 5**: All predictive performance metric results.

**Supplemental Table 6.** 5-fold cross validation results from tests of regularized logistic regression models for binary prediction on the N=80 dataset after regularization strength optimization, with 20 options explored for each model.

| **Penalty** | **Normalization** | **Mean ROC-AUC** |
|---|---|---|
| L1 | Yes | 0.497 |
| L2 | Yes | 0.516 |
| Elastic | Yes | 0.531 |
| L1 | No | 0.475 |
| L2 | No | 0.506 |
| Elastic | No | 0.431 |
